# Broadly inhibitory antibodies against severe malaria virulence proteins

**DOI:** 10.1101/2024.01.25.577124

**Authors:** Raphael A. Reyes, Sai Sundar Rajan Raghavan, Nicholas K. Hurlburt, Viola Introini, Ikhlaq Hussain Kana, Rasmus W. Jensen, Elizabeth Martinez-Scholze, Maria Gestal-Mato, Cristina Bancells Bau, Monica Lisa Fernández-Quintero, Johannes R. Loeffler, James Alexander Ferguson, Wen-Hsin Lee, Greg Michael Martin, Thor G. Theander, Isaac Ssewanyana, Margaret E. Feeney, Bryan Greenhouse, Sebastiaan Bol, Andrew B. Ward, Maria Bernabeu, Marie Pancera, Louise Turner, Evelien M. Bunnik, Thomas Lavstsen

**Affiliations:** Department of Microbiology, Immunology and Molecular Genetics, Long School of Medicine, The University of Texas Health Science Center at San Antonio, San Antonio, TX 78229, USA; Centre for translational Medicine & Parasitology, Department of Immunology and Microbiology, University of Copenhagen and Department of Infectious Diseases, Righospitalet, Copenhagen, Denmark; Department of Integrative Structural and Computational Biology, The Scripps Research Institute, La Jolla, CA 92037, USA; Vaccine and Infectious Disease Division, Fred Hutchinson Cancer Center, Seattle, WA 98109, USA; European Molecular Biology Laboratory (EMBL) Barcelona, Barcelona 08003, Spain; Infectious Disease Research Collaboration, Kampala, Uganda; Department of Medicine, University of California San Francisco, San Francisco, CA 94110, USA; Department of Pediatrics, University of California San Francisco, San Francisco, CA 94110, USA

## Abstract

*Plasmodium falciparum* pathology is driven by the accumulation of parasite-infected erythrocytes in microvessels. This process is mediated by the parasite’s polymorphic erythrocyte membrane protein 1 (PfEMP1) adhesion proteins. A subset of PfEMP1 variants that bind human endothelial protein C receptor (EPCR) through their CIDRα1 domains is responsible for severe malaria pathogenesis. A longstanding question is whether individual antibodies can recognize the large repertoire of circulating PfEMP1 variants. Here, we describe two broadly reactive and binding-inhibitory human monoclonal antibodies against CIDRα1. The antibodies isolated from two different individuals exhibited a similar and consistent EPCR-binding inhibition of 34 CIDRα1 domains, representing five of the six subclasses of CIDRα1. Both antibodies inhibited EPCR binding of both recombinant full-length and native PfEMP1 proteins as well as parasite sequestration in bioengineered 3D brain microvessels under physiologically relevant flow conditions. Structural analyses of the two antibodies in complex with two different CIDRα1 antigen variants reveal similar binding mechanisms that depend on interactions with three highly conserved amino acid residues of the EPCR-binding site in CIDRα1. These broadly reactive antibodies likely represent a common mechanism of acquired immunity to severe malaria and offer novel insights for the design of a vaccine or treatment targeting severe malaria.

## Introduction

Each year, *Plasmodium falciparum* causes approximately 600,000 deaths from malaria, mainly among young children living in sub-Saharan Africa. Ten times as many suffer from severe disease, often with lasting consequences [1]. Malaria pathology is driven by the accumulation of parasite-infected erythrocytes in the microvasculature, resulting in reduced blood flow, inflammation, and endothelial lesions in vital organs [2]. In severe cases, this may lead to organ failure and death. Parasite-infected erythrocytes bind to endothelial cell receptors on the microvasculature via the polymorphic *P. falciparum* erythrocyte membrane proteins (PfEMP1) expressed on their cell surface, thereby avoiding being filtered out by the spleen [3-5]. PfEMP1 are composed of 2 – 10 Duffy binding-like (DBL) and cysteine-rich interdomain region (CIDR) domains [6, 7]. Severe malaria is caused by parasites binding human endothelial protein C receptor (EPCR) through the subset of PfEMP1 that harbor CIDRα1 domains [8-21]. In addition to the adverse microvascular effects caused by sequestration, infected erythrocyte binding to EPCR impairs its normal function, leading to enhanced inflammation and permeability of the microvasculature [22-24]. PfEMP1 are major targets of the humoral immune response to malaria, and antibody reactivity against the CIDRα1 domain family correlates with protection from severe malaria [25-28]. Given their central role in malaria pathogenesis and immunity, the PfEMP1 CIDRα1 domains are attractive targets for a vaccine preventing severe and potentially fatal complications of malaria. However, vaccine development is hampered by the extensive amino acid sequence diversity that has evolved among CIDRα1 domains to escape immune recognition.

EPCR-binding CIDRα1 domains divide into subclasses CIDRα1.1 and CIDRα1.4 – 1.8 and share on average only ∼60% of their 251 amino acids [29]. Structural studies of CIDRα1 domains in complex with EPCR show that despite this extensive sequence diversity, the CIDRα1 fold and the surface chemistry of the EPCR-binding site is conserved in order to retain the capacity to bind to EPCR [6, 30]. We hypothesize that the structural and chemical constraints on the CIDRα1 domain, necessary for its binding to EPCR, may also enable antibody binding to many or even all CIDRα1 variants. However, it is unknown whether such broadly reactive antibodies develop in response to *P. falciparum* infection and how they would interact with the structurally concealed and sequence-diverse CIDRα1 domains. Here, we address these questions through the isolation of broadly inhibitory antibodies to CIDRα1 from two different *P. falciparum*-exposed individuals and structural characterization of their binding mechanism.

## Results

### P. falciparum-exposed individuals develop broadly reactive antibodies against CIDRα1 domains

To identify human monoclonal antibodies (mAbs) against PfEMP1 CIDRα1 domains, we isolated IgG^+^ CIDRα1-specific B cells from three Ugandan adults. B cells were stained using fluorescently labeled recombinant CIDRα1.1 (IT4VAR20 variant) and CIDRα1.4 (HB3VAR03 variant) domains, representing the two most diverse CIDRα1 subclasses (**Figure 1A, B**). B cells with reactivity to either of the domain variants were sorted (**Figure S1** for gating strategy) and cultured individually to differentiate into antibody-secreting cells. The culture supernatants containing mAbs were screened for reactivity against a panel of 34 EPCR-binding CIDRα1 domain variants representing the naturally occurring spectrum of CIDRα1 sequence variants. Using this approach, we isolated seven mAbs with reactivity against CIDRα1, ranging from variant-specific to broadly reactive and able to inhibit the binding of CIDRα1 to EPCR (**Figure 1C**). Three of the mAbs, C7, C62, and C74, showed highly similar reactivity and inhibition patterns, binding to (almost) all CIDRα1.4 – 1.8 variants, but none of the CIDRα1.1 variants (**Figure 1C**). mAbs C7 and C62 were isolated from the same donor. Sequence analysis of the heavy and light chain variable regions revealed that these two mAbs belonged to the same clonal lineage and were nearly identical (**Table S1**). We therefore produced recombinant IgG_1_ versions of mAbs C7 and C74 only. Both were confirmed to bind to and inhibit EPCR binding of diverse CIDRα1 domains as in the initial screen, and in a concentration-dependent manner (**Figure 1D, S2**). Reactivity of both C7 and C74 consistently correlated with inhibition of EPCR binding (**Figure S3**).

**Figure 1:**
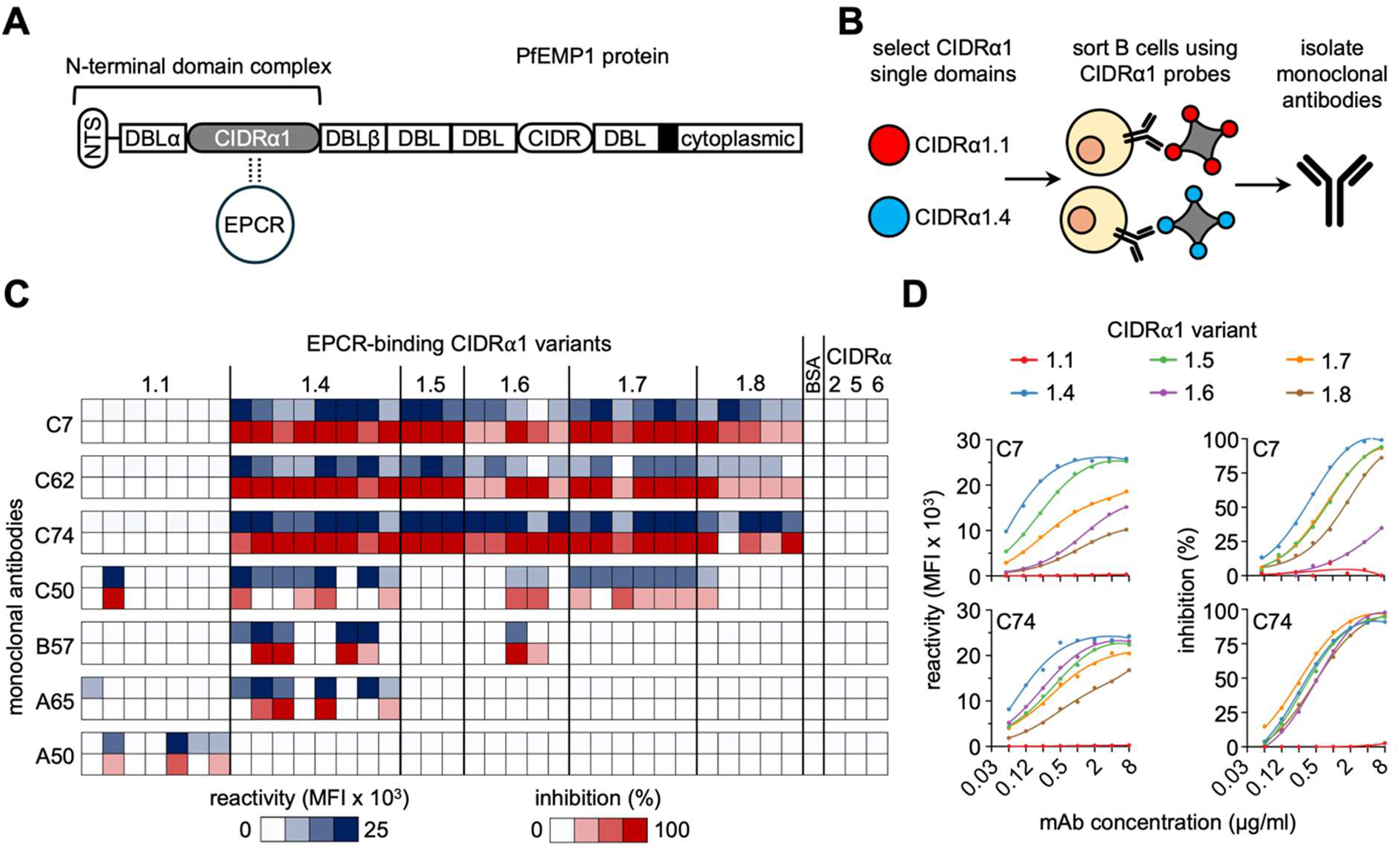
Isolation of monoclonal antibodies against the PfEMP1 CIDRα1 domain. **A)** Schematic representation of CIDRα1-containing multi-domain PfEMP1 proteins with the N-terminal domain complex comprised of the N-terminal segment (NTS), DBLα, and CIDRα1 domain indicated. The EPCR binding site is also shown. **B)** Overview of the experimental strategy to isolate monoclonal antibodies against CIDRα1 domains. **C)** Heatmap showing monoclonal antibody reactivity and inhibition of EPCR binding to a panel of CIDRα1 variants (Luminex assay). Controls include bovine serum albumin (BSA) and CD36-binding CIDRα2, CIDRα5, and CIDRα6 variants. **D)** Titration of monoclonal antibody reactivity and inhibition of EPCR binding to CIDRα1 variants representative of each of the six subclasses for C7 and C74. MFI, median fluorescence intensity.

### Broadly reactive mAbs bind CIDRα1 and multi-domain PfEMP1 with high affinity

In native PfEMP1, the CIDRα1 domain, its flanking DBLα domain, and the N-terminal segment (NTS) comprise the PfEMP1 N-terminal domain complex (**Figure 1A**). The DBLα and CIDRα1 domains interact with each other through central residues of the EPCR binding site on CIDRα1 to form a compact protein structure [30]. As a result, the EPCR-binding site is partially hidden in the unbound PfEMP1. When EPCR binds to CIDRα1, it displaces the DBLα domain and induces a conformational change to CIDRα1 by twisting and turning the EPCR-binding (EB) helix and the EPCR-binding supporting (EBS) helix [30]. Since the mAbs identified here were isolated using single CIDRα1domains, we assessed their ability to bind to a panel of recombinant multi-domain and full-length ectodomain PfEMP1 proteins and inhibit their interaction with EPCR. In line with the results obtained with single CIDRα1 domain variants, C7 and C74 showed reactivity to and EPCR-binding inhibition of PfEMP1 proteins with CIDRα1.4 – 1.8 domains, but not PfEMP1 with CIDRα1.1 domains (**Figure S4A**). Furthermore, biolayer interferometry analyses showed that C7 and C74 Fabs bound CIDRα1 domains with high affinity whether presented individually or within their N-terminal domain complex (**Figure S4B-C)**. Both Fabs exhibited similar on-rate kinetics and very low dissociation rates, mimicking the kinetics of EPCR binding to CIDRα1 [30, 31]. Altogether, these results suggest that C7 and C74 may bind similar epitopes at or near the EPCR binding site in CIDRα1.

### Broadly reactive mAbs bind native PfEMP1 and inhibit endothelial sequestration of parasites

Next, we determined the ability of the C7 and C74 mAbs to bind native PfEMP1 on the surface of *P. falciparum*-infected erythrocytes. For this, mAb binding to parasite-infected erythrocytes predominantly expressing a single PfEMP1 with a CIDRα1.1, CIDRα1.4, or CIDRα1.6 domain was analyzed by flow cytometry (**Figure 2A, S5**). In agreement with binding patterns to the recombinant PfEMP1 proteins, both C7 and C74 IgG_1_ stained parasites expressing CIDRα1.4 and CIDRα1.6 PfEMP1, but not CIDRα1.1 PfEMP1. We then proceeded to test the ability of the mAbs to inhibit the binding of *P. falciparum*-infected erythrocytes to EPCR using CIDRα1.4 PfEMP1-expressing parasites (HB3VAR03). First, we tested binding to recombinant EPCR coated onto plastic (**Figure 2B**). In this static binding assay, EPCR binding was inhibited by the presence of soluble EPCR, C7 IgG_1_, and C74 IgG_1_, but not by a human mAb targeting a non-CIDRα1 PfEMP1.

**Figure 2:**
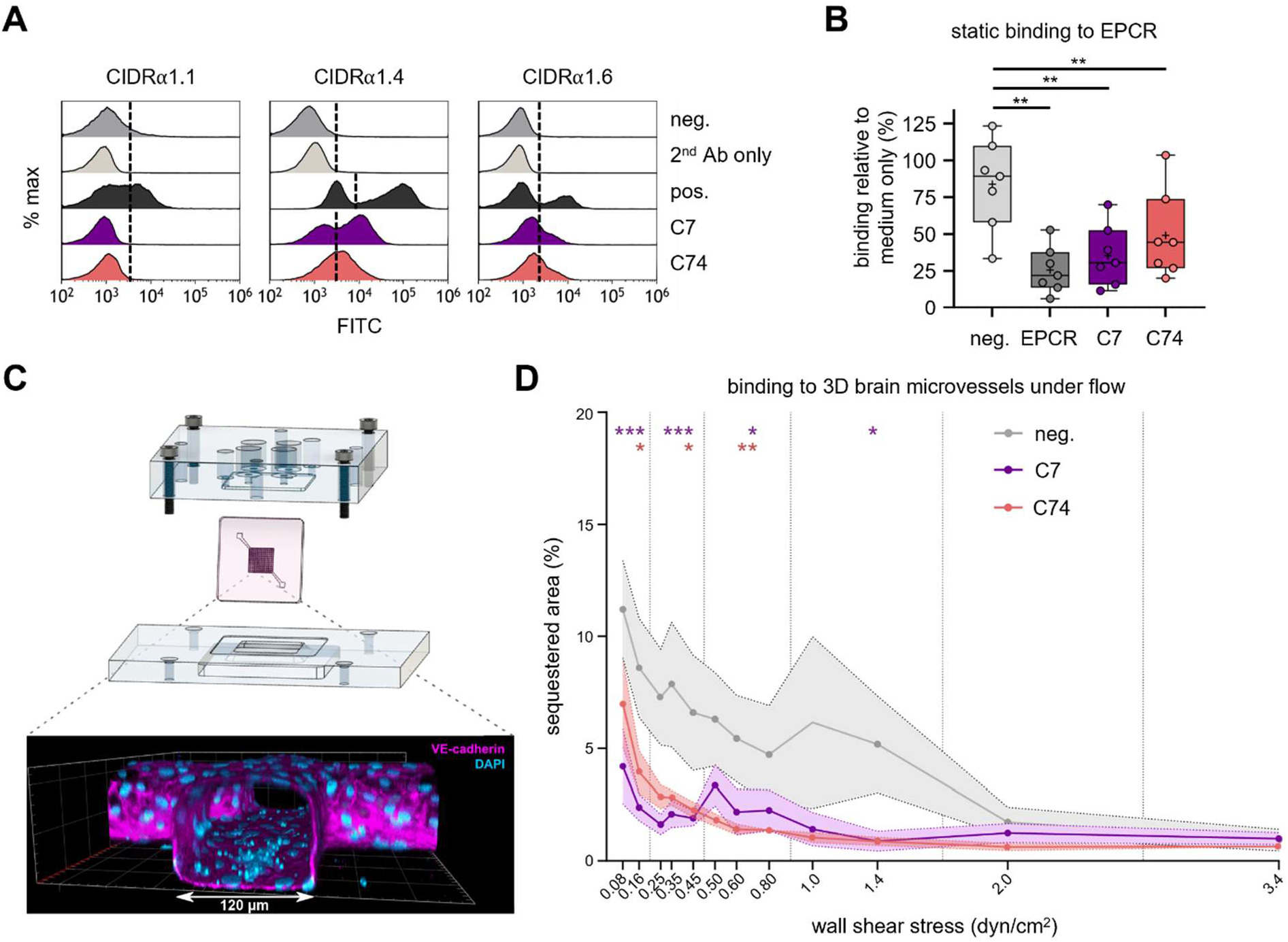
C7 and C74 reactivity to and inhibition of *P. falciparum*-infected erythrocytes. **A)** Flow cytometry analysis showing C7 and C74 (FITC) staining of live *P. falciparum-*infected erythrocytes expressing PfEMP1 variants containing CIDRα1.1 (IT4VAR19), CIDRα1.4 (HB3VAR03), and CIDRα1.6 (IT4VAR18). IgG from rats immunized with the respective PfEMP1 variants were used as positive controls. IgG samples from rats immunized with a heterologous PfEMP1 variant was included as negative controls. Dashed lines indicate the cutoff for positive cells, as determined using non-infected erythrocytes in the same sample. **B)** Binding of *P. falciparum-*infected erythrocytes expressing CIDRα1.4 PfEMP1 (HB3VAR03) to recombinant EPCR under static conditions in culture medium, or in presence of 50 µg/ml C7 or C74. As a negative control, mAb PAM1.4 targeting the non-CIDRα1-containing VAR2CSA PfEMP1 was used. Recombinant soluble EPCR was included as positive control. Binding levels from seven independent experiments are shown. Within each experiment, binding was normalized to the medium only condition that was indexed to 100. A repeated-measures one-way ANOVA followed by Dunnett’s test was used to evaluate differences compared to the negative control. P values shown are from Dunnett’s post-hoc test and are corrected for multiple comparisons. + denotes the mean. **C)** Binding of CIDRα1.4 PfEMP1-expressing *P. falciparum-* infected erythrocytes (HB3VAR03) to human brain endothelial cells in 3D microvessels under flow conditions. Top: schematic of device components used to generate a 13-by-13 3D microfluidic network. Bottom: volumetric reconstruction of a microvessel cross section (120 µm diameter) after immunofluorescence labelling with an anti-human VE-cadherin antibody (magenta) and nuclear staining by DAPI (blue). Parasite nuclei can be identified as smaller, brighter blue foci attached to the bottom endothelial surface (DAPI). **D)** Percentage of endothelial area occupied by sequestered *P. falciparum-*infected erythrocytes in the 3D microvessels at regions exposed to different wall shear stress rates. Dots indicate the median values, and the shaded regions show the interquartile range, from a total of 9, 7, and 9 independent biological replicates for C7 (0.47 mg/ml), C74 (0.4 mg/ml), and the isotype control IgG_1_ (neg., 0.47 mg/ml), respectively. Statistical analyses were performed for binned regions (dotted vertical lines) using a Kruskal-Wallis test, followed by comparisons between IgG_1_ and C7 or C74 using Dunn’s post-hoc test, corrected for multiple comparisons. * P < 0.05, ** P < 0.01, *** P < 0.001.

To assess if C7 and C74 IgG_1_ could also inhibit adhesion of *P. falciparum*-infected erythrocytes to human endothelial cells, we exploited the most physiologically relevant model of *P. falciparum* cytoadherence currently available. In this model, pre-patterned 3D microvessels are seeded with primary human brain microvascular endothelial cells in a microfluidic network to achieve wall shear stress rates found in healthy microvasculature (1 – 3.5 dyn/cm²) and capillaries occluded by *P. falciparum*-infected erythrocytes (0 – 1 dyn/cm²) [32, 33] (**Figure 2C, S6**). Once intact microvessels had formed, the devices were perfused with *P. falciparum*-infected erythrocytes expressing CIDRα1.4 PfEMP1 (HB3VAR03) and sequestration under multiple wall shear stress conditions was quantified. When *P. falciparum*-infected erythrocytes were pre-incubated and perfused with C7 or C74, sequestration was significantly inhibited at all wall shear stress rates below 2 dyn/cm^2^ for C7 and below 1 dyn/cm^2^ for C74, as compared to infected erythrocytes treated with an isotype control IgG (**Figure 2D, S6**).

Collectively, these experiments demonstrate that C7 and C74 IgG_1_ bind native PfEMP1 and inhibit endothelial sequestration of *P. falciparum*-infected erythrocytes by blocking the binding of PfEMP1 to EPCR under physiologically relevant conditions.

### C7 induces an open PfEMP1 conformation and targets the EPCR binding site

To understand the binding mechanism of C7 and C74, we first conducted X-ray crystallography experiments of C7 Fab in complex with the CIDRα1.4 domain (HB3VAR03, **Figure 3A**). This structure, obtained to a resolution of 2.7 Å, showed that C7 Fab bound directly to the EB and EBS helices of the CIDRα1.4 domain (**Figure 3B, Table S3**). Although the CIDRα1.4 domain was only partly resolved, overlay of the CIDRα1.4 domain in complex with EPCR (PDB: 4V3D) [6] and C7 showed that the EB and EBS helices are similarly oriented, perpendicular to the CIDRα1.4 core helices in both interactions (**Figure 3C**). The overlay also demonstrates that C7 directly blocks EPCR binding.

**Figure 3:**
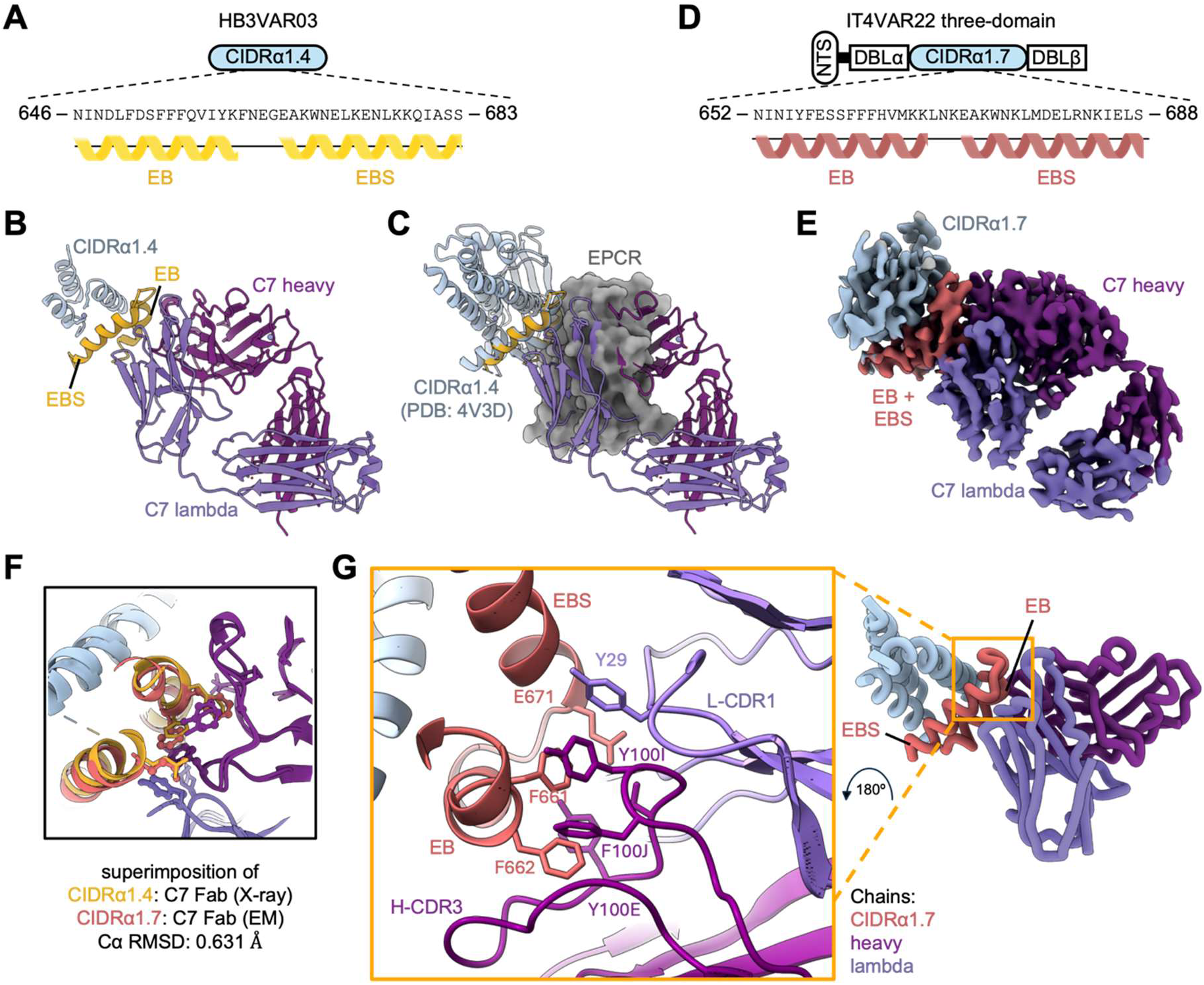
Structural analysis of C7 Fab in complex with recombinant PfEMP1. **A)** Schematic of the single CIDRα1.4 domain (HB3VAR03). EPCR-binding (EB) and supporting (EBS) helices of the CIDRα1.4 domain are shown in yellow. **B)** X-ray crystallography structure of the single CIDRα1.4 domain (HB3VAR03) in complex with C7 Fab shown in cartoon representation. **C)** Overlay of X-ray structure from panel B with the CIDRα1.4 (HB3VAR03):EPCR structure (PDB: 4V3D). EPCR is shown in gray surface representation. **D**) Schematic of the IT4VAR22 three-domain protein. The domain architecture is shown in the top, with EB and EBS helices of CIDRα1.7 highlighted in red. **E)** Cryo-EM map of the IT4VAR22 three domain protein in complex with C7 Fab. **F)** Superimposition of CIDRα1.4 (HB3VAR03) and CIDRα1.7 (IT4VAR22) domains in complex with C7 Fab. The root mean square deviation (RMSD) for alpha-carbon atoms (Cα) in both structures is shown. **G)** Molecular interaction of C7 Fab with EB and EBS helices of CIDRα1.7 (IT4VAR22). π-stacking interactions can be seen between CIDRα1.7 residues F661 and F662 and heavy chain residues Y100E, Y100I, and F100J.

To determine how C7 interacts with CIDRα1 in its more native multi-domain architecture, we performed cryo-electron microscopy (cryo-EM) analysis of C7 Fab in complex with a protein spanning the first three domains of the HB3VAR03 PfEMP1 (NTS-DBLα-CIDRα1.4-DBLβ). We obtained a low-resolution cryo-EM map (∼6 Å) fitted with the HB3VAR03 three-domain structure (PDB: 8C3Y) [30], which confirmed C7’s binding site on CIDRα1 and showed that C7 Fab binding resulted in displacement of the DBLα domain, rendering it flexible **(Figure S7)**. Thus, C7 induces an open conformation of the PfEMP1 N-terminal domain complex similar to that observed following EPCR binding [30].

To further assess the broadly reactive nature of C7, we determined the structure of C7 in complex with the three-domain protein derived from a different PfEMP1 variant (IT4VAR22) containing a CIDRα1.7 domain: NTSA-DBLα-CIDRα1.7-DBLβ (**Figure 3D, Table S4**). The binding interface was resolved at ∼3.2 Å resolution **(Figure 3E, S8 top)**. Here, both the DBLα and DBLβ domain flanking the CIDRα1.7 were unresolved due to flexibility. Superimposing the structures of C7 Fab with the single CIDRα1.4 domain (HB3VAR03) and C7 Fab with the CIDRα1.7 domain (IT4VAR22) (Cα RMSD: 0.631 Å) showed that the C7 epitopes in these two PfEMP1 variants are nearly identical (**Figure 3F**) and mainly targeted CIDRα1 through the heavy chain complementarity determining region (H-CDR3) (**Figure S9**).

The interaction of CIDRα1 with EPCR is centered around a conserved double phenylalanine (FF) motif (F655 and F656 in CIDRα1.4 (HB3VAR03); F661 and F662 in CIDRα1.7 (IT4VAR22)) in the EB helix [6]. This motif establishes a hydrophobic patch that enables the second phenylalanine to protrude into EPCR’s hydrophobic pocket, resulting in a highly stable receptor-ligand complex. The 23 amino acids-long H-CDR3 loop of C7 contains three aromatic residues, Y100E, Y100I, and F100J, which form π-stacking interactions with the central FF motif on the EB helix **(Figure 3G)**. The H-CDR3 forms a hydrophobic grove into which the FF motif protrudes to establish stable binding, mimicking the EPCR binding to CIDRα1 (**Figure S10)**.

In addition to the central FF motif, C7 binding appeared to depend on interactions with glutamic acid residue E671 in CIDRα1.7 (IT4VAR22) and E666 in CIDRα1.4 (HB3VAR03) through H-CDR3 Y100E and S100G (**Figure 3G**). This conserved residue is located at the CIDRα1 EBS helix N-terminal, and stabilizes the EPCR interaction, along with the first aromatic residue of the FF motif [6].

### mAbs C7 and C74 exhibit structurally convergent modes of binding

C7 and C74 displayed similar reactivity and inhibition patterns across the CIDRα1 protein panel, suggesting a similar binding mechanism. To determine how C74 interacts with CIDRα1, we proceeded with cryo-EM analysis of C74 Fab complexed to the CIDRα1.7 (IT4VAR22) three-domain protein. As with the C7 complex, we could not discern DBLα and DBLβ cryo-EM densities, indicating flexibility of these domains induced by C74 binding **(Figure S8)**. However, a ∼3.4 Å resolution map of the C74 Fab-CIDRα1.7 complex was obtained (**Figure 4A, S8 bottom, Table S4**). Similar to C7, the C74 Fab bound the EB and EBS helices through aromatic π-stacking interactions with the FF motif of the EB helix. In C74, this was mediated by residues in both H-CDR2 (Y50 and F58) and H-CDR3 (Y99) (**Figure 4B**). Also similar to C7, mAb C74 contacts E671 through a serine and tyrosine pair in H-CDR2 (S52) and H-CDR3 (Y99). Thus, despite the different H-CDR architectures, the antigen-binding sites formed by the C7 and C74 heavy chains are similar in amino acid composition and mode of binding (**Figure 4C-D**).

**Figure 4:**
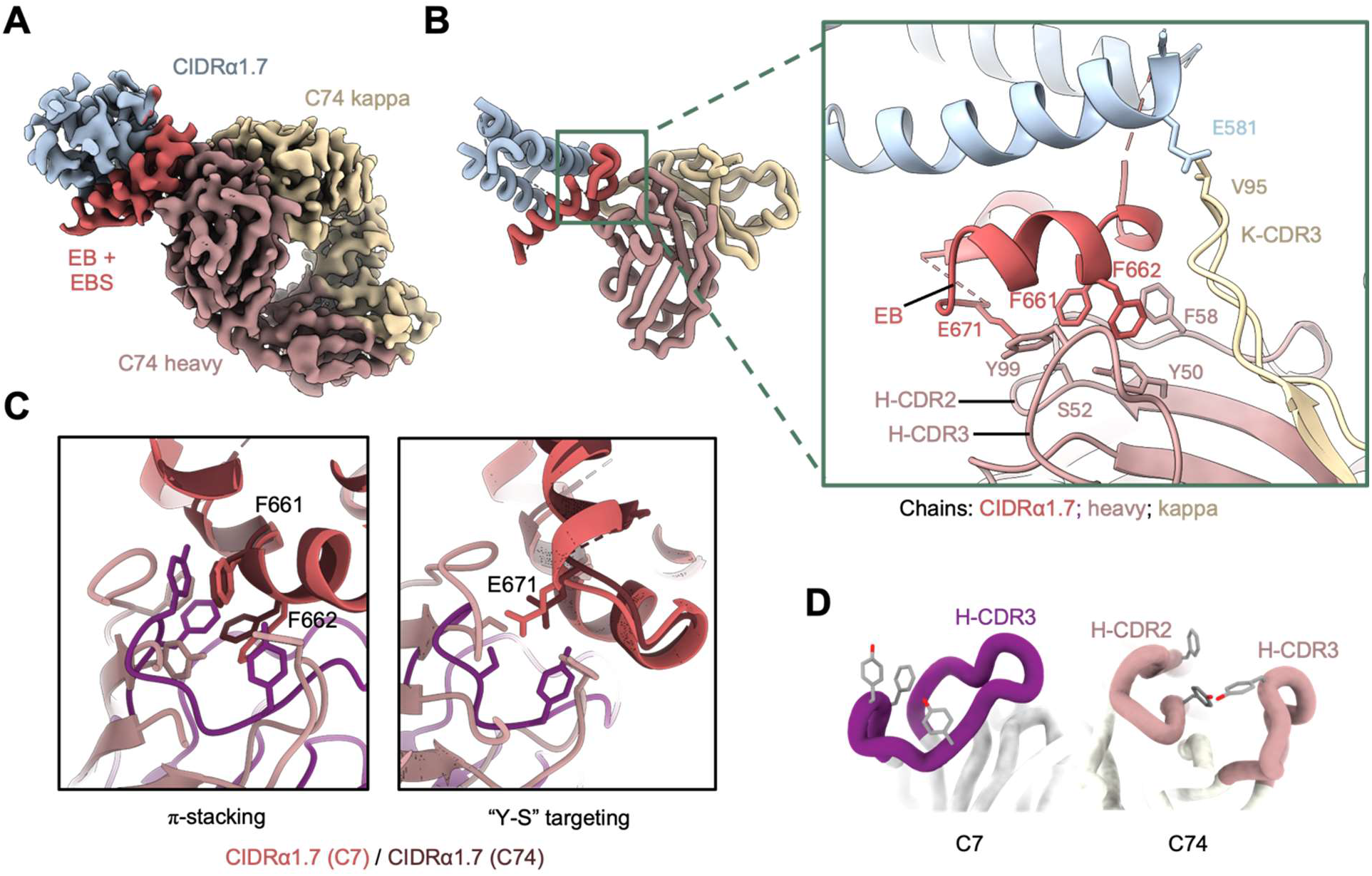
Cryo-EM structure of C74 Fab complexed with the CIDRα1.7 PfEMP1 variant. **A)** Cryo-EM map of C74 Fab complexed with the three-domain protein derived from IT4VAR22 PfEMP1. EPCR binding (EB) and supporting (EBS) helices of CIDRα1.7 are colored in red. **B)** Molecular interaction of C74 Fab with the CIDRα1.7 EB and EBS helices. **C)** Superimposition of aromatic π-stacking interaction with the CIDRα1.7 FF motif (left panel) and superimposition of the Y-S residues of C7 and C74 targeting E671 of IT4VAR22 CIDRα1.7 (right panel). **D)** Key antigen-contacting aromatic residues of C7 and C74 shown in their respective H-CDR conformation.

Both C7 and C74 mirrored the EPCR interaction by inducing the twist-turn of the EB and EBS helices ∼72° clockwise compared to their position in the unbound form **(Figure S11A**). The two mAbs approach the EB and EBS helices with a ∼16° difference in the angle of approach **(Figure S11B**) but are also rotated about 180° in line to the angle of approach. Additionally, in C7, the light chain (L) CDR1 Y29 interacts with the EBS helix glutamic acid residue E671 **(Figure 3G, S9)**, while in C74, L-CDR3 V95 contacts E581 in the core helix 1 of CIDRα1.7 **(Figure 4B, S9)**. E581 is part of a conserved region in CIDRα1.4 – 1.7 forming a secondary EPCR binding site [30]. This site binds EPCR’s β-sheet region through strong hydrogen and hydrophobic contacts to stabilize the conformation of the receptor-binding complex. Overall, despite these differences, the molecular interactions of C7 and C74 are strikingly similar.

### The key C7 and C74 epitope residues are conserved in CIDRα1

The contact residues of C7 and C74 on CIDRα1 largely overlap, with ten shared contact residues on the EB and EBS helices **(Figure 5A)**. Most of these are surface-exposed on the unbound PfEMP1 multi-domain complex (PDB: 8C3Y) [30] (**Figure 5B**). Of the ten shared contact residues, only four residues are conserved. Of these, a tryptophan residue in the EBS helix is found in all CIDRα1 variants. The other three residues, the FF motif in the EB helix and the glutamic acid residue in the EBS helix, are specific to the targeted CIDRα1 variants (**Figure 5C**). Substituting either of the phenylalanine residues in the FF motif or the glutamic acid residue to alanine in the CIDRα1.4 domain (HB3VAR03) resulted in loss of reactivity by both C7 and C74 mAbs, thus validating the relevance of these key epitope residues and their importance for the broad reactivity across the CIDRα1 variants (**Figure 5D**). CIDRα1.1 domains have an LF/Y motif in the EB helix, and a glutamic acid in their N-terminal EBS helix that is shifted one amino acid N-terminal to its position in CIDRα1.4 – 1.8 domains (**Figure 5C**). These differences in the key contact residues match with the mAbs’ reactivity with CIDRα1.4 – 1.8 domains and lack of CIDRα1.1 recognition. Indeed, mutating the CIDRα1.1 sequence to contain an FF motif in the EB helix and a glutamic acid residue in the EBS helix in same position as in CIDRα1.4 – 1.8 domains resulted in gain of reactivity for both C7 and C74 (**Figure 5D**). At the same time, these mutations resulted in a partial loss of EPCR reactivity. Together, these observations demonstrate that C7 and C74 target a conserved epitope and suggest that diversification of this epitope is restricted by the structural and chemical requirements for EPCR binding.

**Figure 5:**
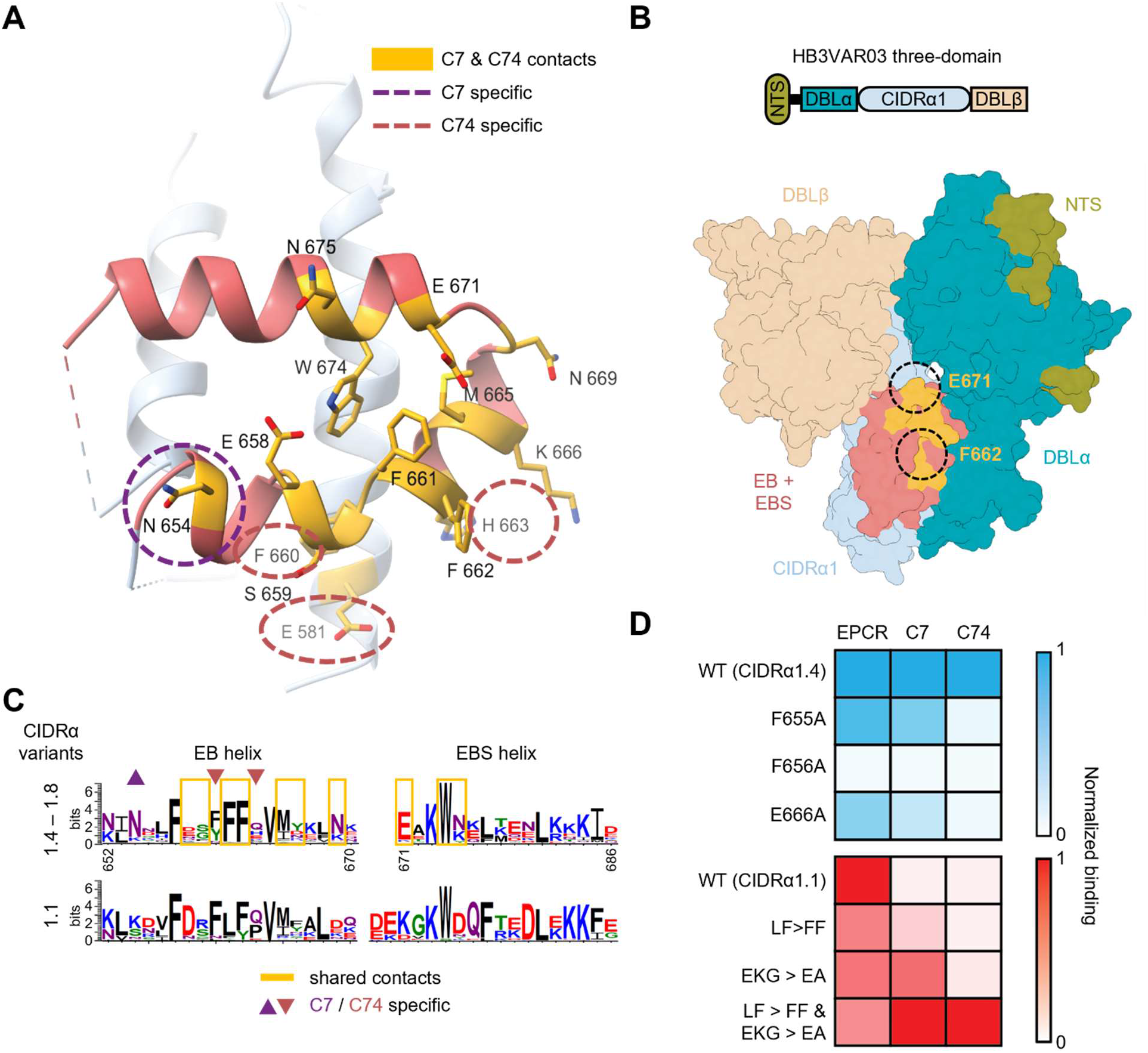
Conservation and surface exposure of C7 and C74 epitope residues. **A)** CIDRα1 residues contacting C7 and C74 mapped onto to the IT4VAR22 CIDRα1.7 domain (circled amino acid residues are mAb specific). **B)** Surface exposure of C7 and C74 contact residues mapped onto the unbound three-domain protein of the CIDRα1.4 PfEMP1, HB3VAR03 (PDB: 8C3Y). Numbering is based on the CIDRα1.7 IT4VAR22 sequence. **C)** Sequence logo plot showing amino acid conservation (colored by chemistry) of the EB and EBS helices in CIDRα1.4 – 1.8 and CIDRα1.1 (numbered relative to the IT4VAR22 sequence). **D**) ELISA reactivity of C7, C74 and recombinant EPCR to the mutated CIDRα1.4 (HB3VAR03) and CIDRα1.1 (IT4VAR20) domains. Wild type OD values >1; data indexed to WT=1.

### Somatic hypermutation is required for broad CIDRα1 reactivity of C7

Although the H-CDR3 length, somatic hypermutations (**Figure S9)**, and light chain pairing are different for C7 and C74 (C7: V_L_3-1; C74: V_K_3-15), both mAbs used heavy chain V gene V_H_3-48. This observation raised the question whether this V gene intrinsically supports broad reactivity to CIDRα1. To answer this, we first generated two inferred-germline (iGL) antibodies: one with the somatic mutations of only the C7 heavy chain (V and J gene segments) reverted to germline (iGL_H_) and one in which mutations in both heavy and light chains (V and J gene segments) were reverted (iGL_HL_). In both antibodies, the untemplated CDR3s were left unchanged (**Figure 6A**). Both antibodies showed loss of broad reactivity, with the iGL_H_ and iGL_HL_ antibodies binding to only seven and three out of 20 CIDRα1 variants bound by C7, respectively **(Figure 6B)**. Assessment of the binding kinetics of iGL_H_ and iGL_HL_ to the IT4VAR22 three-domain protein using BLI showed a markedly reduced affinity due to both lower on-rates and higher off-rates **(Figure S12A)**. Thus, despite the presence of the intact H-CDR3, the germline-reverted antibodies showed reduced reactivity to CIDRα1. This result is not unexpected since mutated residues outside H-CDR3 also contact CIDRα1 (**Figure S9**).

**Figure 6:**
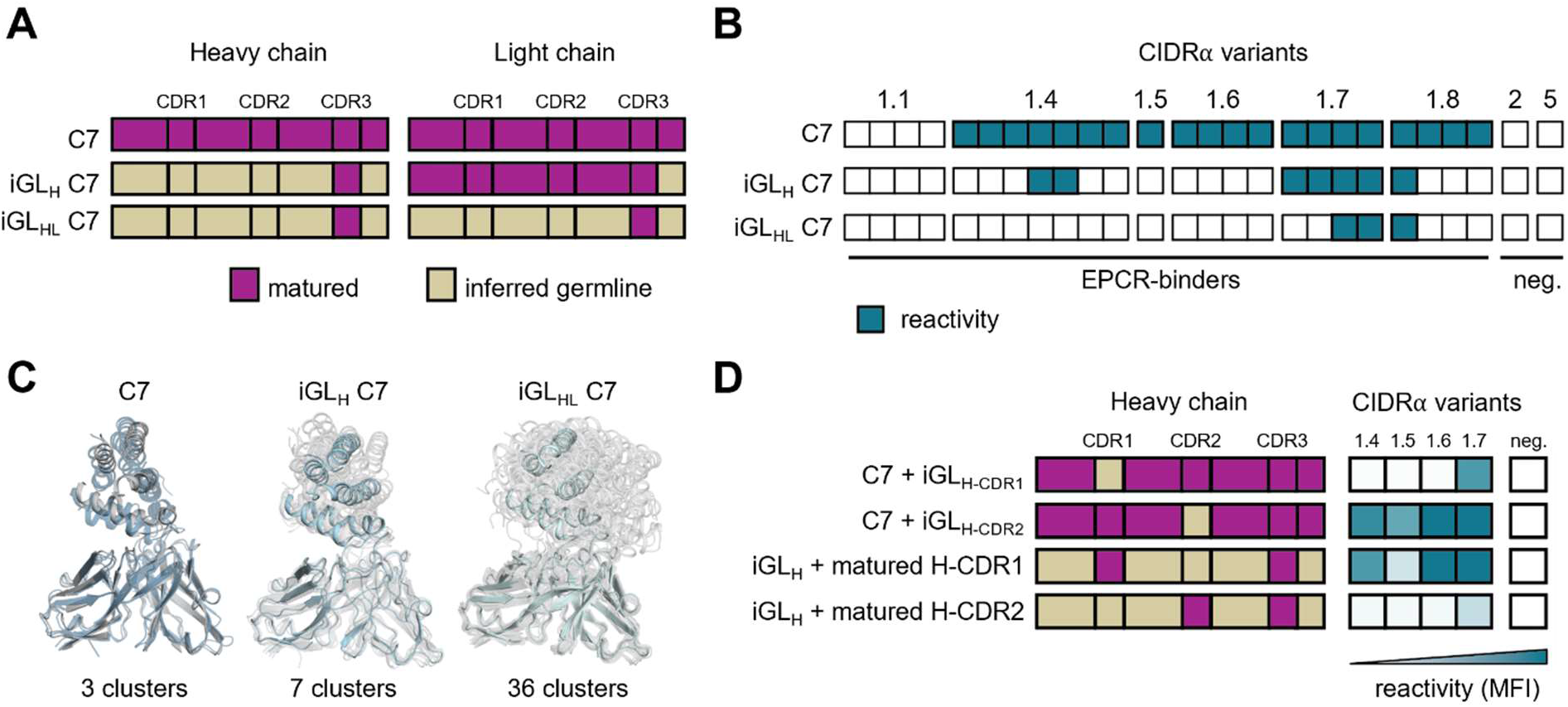
Analysis of C7 germline mAbs. **A)** Depiction of inferred germline antibody designs, showing mAb C7 (top), C7 containing an inferred germline heavy chain variable region (iGL_H_, middle), and C7 containing inferred heavy and light chain variable regions (iGL_HL_, bottom). **B)** Reactivity of C7, iGL_H_ C7 and iGL_HL_ C7 across a panel of recombinant CIDRα1 protein domains and negative control (neg.) CD36-binding CIDRα2/5 domains. Blue squares indicate positive reactivity. **C)** All-atom molecular dynamics simulation of the antigen-bound structure of C7, iGL_H_ C7 and iGL_HL_ C7 with the number of distinct structural clusters indicated. **D)** Reactivity of C7 with germline-reverted H-CDR1 or H-CDR2 as well as iGL_H_ C7 with matured H-CDR1 or H-CDR2 to select representative CIDRα1 protein domains and a negative control (neg.).

To further assess the molecular basis of this reduced reactivity, we conducted all-atomistic molecular dynamics simulations on the free variable regions of C7 to explore the free energy landscape of the H-CDR3 loop in the mutated and germline-reverted C7 antibodies (**Figure S12B**). Here, the H-CDR3 of C7 exhibited a single dominant conformation, whereas iGL_H_ and iGL_HL_ C7 displayed high conformational diversity of the H-CDR3 loop (**Figure S12C**). Accordingly, models of C7, iGL_H_ C7 and iGL_HL_ C7 in complex with CIDRα1.7 (IT4VAR22) showed destabilization of the germline-reverted antibody-antigen complexes (**Figure 6C**), with a reduction in the overall interaction profile of the key H-CDR3 aromatic residues in the germline-reverted antibodies (**Table S2**).

The structure of C7 Fab suggested that stabilization of the long H-CDR3 loop was particularly dependent on H32 and G33 in H-CDR1 as well as K56 in H-CDR2 **(Figure S12D)**. To assess the importance of H-CDR1 and H-CDR2 in supporting broad antigen reactivity by H-CDR3, we generated versions of C7 with only H-CDR1 or H-CDR2 reverted to germline. Reversion of H-CDR1 but not H-CDR2 caused loss of broad reactivity (**Figure 6D**). In agreement with this, introduction of the mutated H-CDR1 but not H-CDR2 into iGL_H_ C7 restored broad reactivity. Altogether, these data demonstrate the importance of the matured H-CDR1 in facilitating broad reactivity of C7 and underscore the importance of somatic hypermutation for acquiring breadth of reactivity against the CIDRα1 protein family.

## Discussion

Studies in the 1980s [34-37] identified the variant PfEMP1 adhesion antigens as the main targets of exposure-dependent immunity to malaria [38]. Ever since, the molecular mechanism and cross-reactive potential of anti-PfEMP1 antibodies have been subject of study and debate [39-42]. The identification of PfEMP1’s interaction with EPCR as the common trait of parasites causing severe malaria [6, 8-21, 24] has reinforced the notion that individuals exposed to *P. falciparum* infection may develop broadly reactive antibodies capable of inhibiting this central driver of severe malaria pathogenesis. The functional and molecular characterization of mAbs C7 and C74 presented in this study not only provides conclusive evidence for the presence of broadly inhibitory antibodies against EPCR-binding PfEMP1, but also unveils that such antibodies likely share a uniform mode of binding.

Molecular resolution of C7 and C74 interactions with PfEMP1’s CIDRα1 domain revealed similar structural and chemical characteristics of their heavy chain paratopes and epitopes. Central to the binding mechanism of both mAbs is the interaction with the conserved phenylalanine and glutamic acid residues of the EPCR binding (EB) and EPCR-binding supporting (EBS) helices in CIDRα1.4 – 1.8. The mAbs displayed structurally homologous hydrophobic pockets formed by aromatic residues to bind the central EB helix FF motif. Similarly, both antibodies presented the same arrangement of serine and tyrosine residues to contact the glutamic acid conserved in the EBS helix of CIDRα1.4 – 1.8 domains. This observation explains the nearly identical pattern of reactivity and inhibition across CIDRα1 variants by mAbs C7 and C74, despite their different sequence composition. Although C7 and C74 share heavy chain V gene allele V_H_3-48, their similar mode of binding is supported by different structural features. In C74, broad reactivity depended on direct interactions of residues in H-CDR2 and H-CDR3 with CIDRα1, while for C7, broad reactivity relied on its relatively long H-CDR3 loop. Additionally, the conformation of C7’s H-CDR3 was dependent on interactions with the mutated H-CDR1.

These observations are important for several reasons. Firstly, they suggest that the C7/C74-like binding mechanism is commonly applied by broadly inhibitory antibodies against EPCR-binding PfEMP1, and that such antibodies can likely develop through different precursors and maturation routes. However, further studies will need to be done to determine if the common V_H_-3-48 precursor found in these two mAbs is indeed a stochastic event. Analyses of the inferred germline antibodies suggest that somatic mutations are required to gain broad reactivity. However, the degree of *P. falciparum*-exposure and the number of antigen variants required for development of broadly inhibitory antibodies is unknown and may not depend on the specific order in which CIDRα1 variants are encountered. Secondly, it suggests that CIDRα1.4 – 1.8 PfEMP1 cannot vary the targeted, conserved residues without compromising receptor binding or resembling another common serotype. In support of this, both the FF motif in the EB helix and the glutamic acid residue in the EBS helix are diversified in CIDRα1.1 PfEMP1 that can bind EPCR, yet is not recognized by C7 and C74 [7]. When these key amino acid residues were introduced into the CIDRα1.1 domain, they conferred antibody reactivity at the cost of reduced EPCR binding. The notion of two major CIDRα1 serotypes aligns with serological studies of CIDRα1 reactivity, although such data also suggest the co-existence of fully CIDRα1 pan-reactive antibodies [6]. Further studies are needed to elucidate the inhibitory potential and binding mechanism of such antibodies.

Models reflecting the impact of parasite sequestration on severe malaria pathogenesis are difficult to establish as PfEMP1 proteins are unique to *Plasmodium* parasites infecting humans and great apes, and highly adapted to their endothelial receptors [43, 44]. However, the link between EPCR-binding PfEMP1 and severe malaria is well established [8-21]. As PfEMP1 are multi-domain proteins known to bind multiple human receptors, we tested the ability of mAbs C7 and C74 to inhibit binding of *P. falciparum-*infected erythrocytes to recombinant EPCR and endothelial cells under *in vitro* conditions mimicking those present in capillaries and post-capillary venules [32, 33]. The observation that both antibodies inhibited parasite cytoadherence under diverse flow conditions suggests that these antibodies would also affect tissue sequestration *in vivo*.

The data presented here provide evidence that *P. falciparum*-exposed humans develop broadly reactive and inhibitory monoclonal antibodies against severe malaria-associated CIDRα1 PfEMP1. The observation that two antibodies from different individuals exhibit similar modes of binding suggests this interaction represents a common solution of the acquired humoral immune response against severe malaria-associated PfEMP1. This will allow the exploration of such antibodies for antibody therapies aimed at preventing severe malaria. In addition, structural data identified the minimal epitope required for broad antibody recognition. This epitope with low structural complexity comprising a helix-turn-helix motif is highly amenable to epitope scaffold vaccine design.

## Acknowledgements

The authors thank H.L. Turner for cryo-EM technical support, C.A. Bowman for computing support, and W. Lessin for the maintenance and administration of the cryo-EM facility at The Scripps Research Institute. We thank J. Copps for cell culture maintenance and assisting in recombinant protein production. We thank L.H. Fuhrmann for all the laboratory and administrative management. We thank the J. B. Pendleton Charitable Trust for its generous support of Formulatrix robotic instruments. X-ray diffraction data was collected at the Berkeley Center for Structural Biology beamline 5.0.2, which is supported in part by the Howard Hughes Medical Institute. The Advanced Light Source is supported by the Director, Office of Science, Office of Basic Energy Sciences, of the United States Department of Energy under contract number DE-AC02-05CH11231. We thank Lea Barfod for the use of PAM1.4. SSRR, LT, and TL were funded by The Lundbeck Foundation (R344-2020-934) and the Independent Research Fund Denmark (9039-00285A). VI and MB work was supported by a Marie Skłodowska-Curie Actions post-doctoral fellowship FEBBRIS [101026717]) to VI and by core program funding of the European Molecular Biology Laboratory (EMBL). This work was supported by the National Institutes of Health (R01 AI153425 to EMB, F31 AI169993 to RAR, TL1 TR002647 to RAR, R01 AI093615 to MEF, U19 AI150741 to BG, and U19 AI089674). Data were generated in the Flow Cytometry Shared Resource Facility, which is supported by UT Health San Antonio, NIH-NCI P30 CA054174 (Mays Cancer Center at UT Health), CPRIT (RP210126) and NIH Shared Instrument grant 1S10OD030432-01 (S10 grant towards the purchase of a DB FACSAria Fusion). The 3T3-msCD40L cell line was a kind gift from Dr. Mark Connors (National Institute of Allergy and Infectious Diseases). The plasmid encoding rat CD200 was a kind gift from Dr. Gavin Wright (Wellcome Sanger Institute; Addgene plasmid #36152). We acknowledge CHRONOS for awarding us access to Piz Daint at CSCS, Switzerland.

## Author Contributions

Conceptualization: R.A.R., S.S.R.R., N.K.H., M.B., M.P., A.B.W., L.T., E.M.B., and T.L.; Methodology: R.A.R., S.S.R.R., N.K.H., J.A.F., M.L.F.Q., J.R.L., I.K.H., M.B., M.P., A.B.W., L.T., E.M.B., and T.L.; Investigation: R.A.R., S.S.R.R., N.K.H., R.W.J, V.I., M.G.M., E.M.S., S.B., M.P., L.T., E.M.B., and T.L.; Validation: R.A.R., S.S.R.R., N.K.H., J.A.F., I.K.H., M.L.F.Q., C.B.B., J.R.L., W.L., V.I., G.M.M., A.B.W., M.P., E.M.B., and T.L; Formal analysis: R.A.R., S.S.R.R., N.K.H., I.K.H., J.A.F., V.I., M.L.F.Q., J.R.L., S.B., M.B., L.T., and E.M.B.; Data Curation: R.A.R., S.S.R.R., N.K.H., R.W.J., M.G., M.L.F., G.M.M., A.B.W., L.T., E.M.B., and T.L.; Resources: I.S., M.E.F., B.G., M.B., M.P., A.B.W., T.G.T., L.T., E.M.B., and T.L.; Writing-Original draft: R.A.R., S.S.R.R., E.M.B., and T.L.; Writing-Review & editing: N.K.H, V.I., M.P., M.B., S.B.; Visualization: R.A.R., S.S.R.R., N.K.H., S.B., M.B., E.M.B., and T.L.; Supervision: S.B., M.P., M.B., A.B.W., L.T., E.M.B., and T.L.; Project Administration: E.M.B., and T.L.; Funding Acquisition: T.G.T., A.W.B., M.B., M.P., E.M.B., and T.L. All authors contributed to the article and approved the submitted version.

## Methods

### Study design and ethics approval

Individuals included in this study were residents of the Nagongera sub-county in Tororo District, Uganda. This region was historically characterized by extremely high malaria transmission intensity, with an estimated annual entomological inoculation rate of 125 infectious bites per person per year [45]. Individuals included in this study were enrolled in the Program for Resistance, Immunology, Surveillance, and Modeling of Malaria (PRISM) program [46] and have provided written consent for the use of their samples for research. The PRISM cohort study was approved by the Makerere University School of Medicine Research and Ethics Committee (SOMREC) and the University of California, San Francisco Human Research Protection Program. The use of cohort samples for this study was approved by the Institutional Review Board of the University of Texas Health Science Center at San Antonio. Donor 2 was an anonymous blood donor at Mbale regional blood bank in Eastern Uganda, who consented to the use of their blood for research. The use of samples from anonymous blood donors was not considered human research by the Institutional Review Board of the University of Texas Health Science Center at San Antonio due to the lack of any identifiable information and was therefore exempt from review.

### Isolation of CIDRα1 specific B cells

Fluorophore-conjugated antigen tetramers were generated from recombinant HB3VAR03 CIDRα1.4 and IT4VAR20 CIDRα1.1 domains fused with C-terminal StrepTagII. These proteins were produced in baculovirus-infected insect cells as previously described [6]. Antigen tetramers were synthesized by incubating CIDRα1 protein with fluorophore-conjugated streptavidin overnight at 4°C at a molar ratio of 6:1 with rotation. To make decoy tetramers, streptavidin-PE or streptavidin-BV421 was first conjugated to Alexa-fluor 647 (Thermo #A20186) per manufacturer’s instructions. This double-conjugated streptavidin was then coupled to *R. norvegicus* CD200 (Addgene #36152) as described for CIDRα1.

Cryopreserved PBMCs were thawed and DMSO was washed out by mixing with pre-warmed thawing medium (IMDM GlutaMAX supplemented with 10% heat-inactivated FBS (USA origin) and 0.01% Universal Nuclease (Thermo #88700). After centrifugation (250 × g, 5 min.), the cell pellet was resuspended in thawing medium and the cells were counted. Next, cells were centrifuged (250 × g, 5 min.) and resuspended in PBS supplemented with heat-inactivated FBS (2% f/c) and EDTA (1 mM f/c) at 50 million live cells/mL and filtered through a 35 µm sterile filter cap (Corning #352235) to break apart aggregated cells. B cells were isolated using the EasySep Human B Cell Isolation Kit (StemCell #17954) or the MojoSort Human Pan B Cell Isolation Kit (BioLegend #480082) according to the manufacturer’s instructions. B cells were washed with PBS, centrifuged (250 × g, 5 min.), resuspended in 1 mL PBS containing 1 µL live/dead stain (LIVE/DEAD Fixable Aqua Dead Cell Stain Kit (Thermo #L34965)) and incubated on ice for 30 min. Cells were subsequently washed with cold PBS containing 1% BSA (PBS/BSA) (250 × g, 5 min., 4⁰C), resuspended in a 100 µL PBS/BSA and25 µM of each antigen and decoy tetramers, and incubated at 4°C for 30 min. Next, the cells were washed with cold PBS/BSA (250 × g, 5 min., 4⁰C) and incubated at 4°C for 30 min. with 100 µL B cell surface marker antibody cocktail (Table 1B) diluted in PBS/BSA. Finally, the cells were washed with 3 mL cold PBS/BSA, resuspended in cold PBS/BSA to 20 – 30 million cells/mL and filtered into a FACS tube through a 35 μm sterile filter cap. Cells were immediately run on a FACSAria III and antigen-specific B cells were sorted into IMDM GlutaMAX/ 10% FBS.

### Monoclonal B cell activation and expansion

One day prior to sorting B cells, wells of a 96-well plate were each seeded with 30,000 adherent, CD40L-expressing 3T3 cells (kind gift from Dr. Mark Connors, NIH) in 100 µL IMDM GlutaMAX/ 10% FBS containing 2× MycoZap Plus-PR (Lonza #VZA-2021), 100 ng/mL human IL-2 (GoldBio #1110-02-50), and 100 ng/mL human IL-21 (GoldBio #1110-21-10) to promote expansion and differentiation of B cells into antibody-secreting cells. Plates were incubated O/N at 37°C and 8% CO_2_. Immediately after sorting, B cells were resuspended at a concentration of 1 cell per 100 µL IMDM GlutaMAX/10% FBS, added to the 100 µL culture media with supplements already present in the plates to a total volume of 200 µL, and incubated at 37°C and 8% CO_2_. Fourteen days later, the IgG concentration in the supernatant was determined by an enzyme-linked immunosorbent assay (ELISA).

### Recombinant protein production

The bead-coupled panel of CIDRα1 domains has been described in [6], the full-length ectodomain of IT4VAR20 is described in [47], and the three-domain protein of HB3VAR03 is described in [30]. These proteins, as well as the single domain CIDRα1 mutants and multi-domain PfEMP1 proteins used here were produced as His-tagged proteins using the same methods as described in [29] with the following borders: full-length ectodomain of HB3VAR03: amino acids 1-2746, three-domain protein of IT4VAR22: amino acids 1-1153, N-terminal domain complex of IT4VAR07: amino acids 1-713, three-domain protein of IT4VAR19: amino acids 1-1221. Mutated variants of CIDRα1.4 HB3VAR03 were derived from HB3VAR03 amino acids 499-719, and mutated variants of CIDRα1.1 IT4VAR20 were derived from IT4VAR20 amino acids 503-724 (all with a C556S substitution to remove a free cysteine),

DNA sequences encoding heavy and light chains of sequenced monoclonal antibodies were codon-optimized and synthesized for recombinant antibody production in Expi293F cells as previously described [48, 49]. Germline heavy chain variable region gene segments were identified using the International Immunological Information System (IMGT) gene database and the VQuest sequence alignment tool. Germline sequences for C7 variable regions were synthesized as lyophilized double-stranded DNA gBlock fragments (IDT, in 250 – 500 ng quantities). The gBlocks contained *EcoRI* and *NheI* restriction sites, allowing for cloning into the IgG_1_ heavy chain expression vector.

Recombinant antibodies were purified using protein G magnetic beads (Promega #G7472), HiTrap protein G HP columns (Cytiva #17040401), or HiTrap MabSelect PrismA protein A columns (Cytiva #17549852). For production of antibody Fabs, genes encoding the variable regions of C7 and C74 heavy chains were cloned into a pMN vector with a human CH1 domain and a C-terminal His-tag using In-Fusion cloning system (Takara Bio #639649). The full light chains of C7 and C74 were cloned into the pMN vector without any purification tag using the same method. The Fab was expressed in HEK293E cells use PEI as transfection reagent. Cultures were harvested after 7 days and Fabs were purified from culture supernatant using His60 Ni Superflow resin (Takara Bio #635660). Fabs were eluted from the column using a buffer of 50 mM Tris, 300 mM NaCl, pH 7.5 with 150 mM imidazole. Both Fabs were further purified using a Superdex 200 16/600 size exclusion column (Cytiva #28-9893-35) using an AKTApure system (GE Biosciences) and buffer exchanged into 5 mM HEPES, 150 mM NaCl, pH 7.5. C7 Fab was mixed with CIDRα1.4 HB3VAR03 at a 1.5:1 Fab-to-CIDRα1.4 molar ratio and the complex was purified using a Superdex 200 16/600 size exclusion column (Cytiva) and concentrated to an OD_280_=10.

### Enzyme-linked immunosorbent, Luminex and biolayer interferometry assays

To detect IgG, 96-well ELISA plates (Corning #3361) were coated with goat anti-human IgG (Sigma #I2136) antibody at a concentration of 4 µg/mL in PBS, at a volume of 100 µL per well. After a one-hour incubation at 37°C or O/N at 4°C, each well was washed once using slowly running (approximately 900 mL/min.) deionized water. All subsequent washes were performed this way. One hundred fifty µL blocking buffer (one-third Non-Animal Protein (NAP)-Blocker (G-Biosciences #786-190P) and two-thirds PBS) was added to each well to prevent non-specific binding. After one hour of incubation at 37°C, the wells were washed three times, after which 5 µL B cell culture supernatant diluted 20× with dilution buffer (1% NAP Blocker in PBS) to a total volume of 100 µL was added per well. Plates were incubated for two hours at 37°C and washed five times. Then, 100 µL 1:2500 diluted (1% NAP Blocker in PBS) HRP-conjugated anti-human IgG antibody (BioLegend #410902) was added to each well. After incubation for one hour at 37°C and three washes, HRP activity was detected using 50 µL TMB (Thermo #PI34024). Plates were incubated in the dark at RT and the oxidation reaction was stopped by adding 50 µL 0.18 M H_2_SO_4_ per well when the negative controls (buffer only, no culture supernatant) started to color. Absorbance was measured at 450 nm using a BioTek Synergy H4 microplate reader. A human IgG (Sigma #I2511) standard curve (10 three-fold serial dilutions starting at 20 µg/mL) was used to quantify samples.

Monoclonal antibody reactivity and EPCR binding inhibition of recombinant PfEMP1 proteins determined by ELISA were performed as described previously in [50]. In brief, wells were coated with 50 μL 0.2 μM recombinant PfEMP1 or EPCR. Monoclonal antibodies at a 0.4 mg/mL concentration were added at a 1:80 dilution and reactivity was detected using rabbit anti-human IgG HRP at 1:3000. For inhibition assays, recombinant PfEMP1 proteins were added to coated EPCR and detected with and without pre-incubation with monoclonal antibody using anti-His HRP antibodies. Monoclonal antibody reactivity and EPCR binding inhibition to a panel of recombinant single CIDRα1 proteins [50] was determined using a Luminex assay as described previously [51].

Fab binding kinetics were measured using biolayer interferometry on an Octet 96 Red instrument (ForteBio) using FAB2G biosensors (Sartorius). C7 and C74 Fabs were diluted to 10 μg/mL in Kinetics Buffer (KB: 1× PBS, 0.01% Tween 20, 0.01% BSA, and 0.005% NaN_3_, pH 7.4). Each analyte was serially diluted to desired concentration in KB. Fabs were loaded onto biosensor until a threshold of 1.0 nm shift was reached. After loading, biosensors were placed in KB to 60 s for a baseline reading. Biosensors were then immersed in analyte for 600 – 900 s association phase, followed by a 300 – 600 s dissociation phase in KB. The background signal from a biosensor loaded with Fab but with no analyte was subtracted from each loaded biosensor. Kinetic analyses were performed at least twice with independently prepared analyte dilution series. Curve fitting was performed using a 1:1 binding model using the Octet data analysis software, version 9. Mean K_on_ and K_off_ values were determined by averaging all binding curves that had a R^2^ value >0.95.

### Plasmodium falciparum culture

*P. falciparum* clones HB3 and IT4 were maintained and selected for expression of PfEMP1 variants HB3VAR03, IT4VAR18, or IT4VAR19 as previously described in [50, 52]. Briefly, parasites were maintained at 4% hematocrit in O+ human erythrocytes in parasite growth medium (RPMI-1640 containing 25 mM HEPES, 4 mM L-glutamine, 5 g/L AlbuMAX II, 0.02 g/L hypoxanthine, and 25 µg/mL gentamicin). Enrichment of parasites expressing specific PfEMP1 variants was done by selection for binding to human recombinant EPCR or antibodies raised against recombinant PfEMP1 constructs (IgG purified from rats immunized with full-length HB3VAR03 protein; polyclonal rat anti-serum against IT4VAR19 CIDRα1.1 or IT4VAR18 CIDRα1.6). PfEMP1 expression phenotypes were assessed by reverse transcription quantitative PCR (qPCR) using primers specific for each *var* gene of HB3 or IT4 [53] and by flow cytometry using the antibodies mentioned above. For flow cytometry analysis of parasite-infected erythrocytes, mature trophozoite/early schizont stage parasites were purified with a magnetic separation unit and stained with 20 μM Hoechst 33342 (Thermo #62249) for 10 min. in RPMI. After three washes, 0.3 × 10^6^ parasites, re-suspended in 100 μL PBS with 2% BSA, were transferred into the wells of a 96-well U-bottom plate and incubated with an optimized dilution/concentration of control plasma/IgG or with C7/C74 IgG (100 μg/mL for the HB3VAR03- and IT4VAR19-expressing parasite lines, and 50 μg/mL for the IT4VAR18 parasite line) for one hour. The plate was centrifuged at 500 × g for 5 min., supernatants were discarded, and pellets were re-suspended in Alexa Fluor 488 goat anti-rat (Invitrogen #A11006) or anti-human (Invitrogen #A11013) secondary antibodies at 20 μg/mL in PBS/2% BSA for 30 min. Following two washes with PBS/2% BSA, samples were analyzed using a CytoFLEX S flow cytometer (Beckman Coulter Life Sciences). Hoechst 33342 and Alexa Fluor 488 staining were read in BV421 and FITC channels, respectively. Flow cytometry data were analyzed with Kaluza Analysis Software version 2.1 (Beckman Coulter Life Sciences). HB3VAR03-expressing parasites used for 3D microvessel cytoadhesion assays were cultured using human O+ erythrocytes in RPMI-1640 medium supplemented with 10% human type B+ serum at 37°C, 90% N_2_, 5% CO_2_, and 5% O_2_. Parasites used for these experiments underwent between 18 to 31 replication cycles in culture after panning on primary human brain microvascular endothelial cells (HBMECs).

### Antibody inhibition of binding to EPCR in static assay

The static receptor-binding assays were performed as previously described [47]. Briefly, 20 μL aliquots of 20 μg/mL recombinant EPCR were spotted on Petri dishes (Falcon #351029) with spots distributed radially at an equal distance from the center. The dishes were incubated for 2 hours at 37°C in a humidified box. Unbound EPCR protein was aspirated, and spots were blocked with PBS/2% BSA for at least 30 min. at 37°C. Late-stage HB3VAR03-expressing trophozoites (3 × 10^6^/mL) were pre-incubated with negative control IgG (anti-VAR2CSA PAM1.4), mAbs C7 or C74, or recombinant EPCR (all at 50 μg/mL in RPMI) or RPMI only for 15 min. After removal of the block droplet, 20 µl of parasite suspension was added to designated spots. Following an hour incubation, parasite suspension droplets were aspirated, and the dishes were washed by adding 20 mL PBS/2% BSA and incubating for 20 min. on a tilting table at 20 rpm. Adhered infected erythrocytes were fixed with 1.5 % v/v glutaraldehyde for 10 min. and stained with Giemsa for 20 min. Images (five per spot) were captured by light microscopy using 10× objective. Parasites were counted using Fiji software (ImageJ 2.9.0). Percent inhibition was calculated relative to binding in the presence of the negative control antibody.

### 3D brain microvessel fabrication

Primary HBMECs (Cell Systems #ACBRI 376) were grown in a flask coated with poly-L-lysine (Sigma #P8920) up to passage 9 before they were seeded in microvessels. 3D brain microvessel devices were prepared as described previously in [32, 54]. The top part of the microvessels is generated by injecting type I collagen (7.5 mg/mL) into the space created between the top plexiglass jig and a polydimethylsiloxane (PDMS) mold with a 13-by-13 grid geometry, fabricated using soft lithography as shown in **Figure 2** and **Figure S6**. The bottom part consists of a flat layer of collagen, compressed between a flat PDMS stamp and a 22-by-22 mm coverslip positioned on the bottom jig. After 30 min. of gelation at 37°C, the PDMS stamps were removed, and the top and bottom jigs were sealed creating a three-dimensional network in the collagen. An hour later 8 μL primary HBMECs at a concentration of 7 × 10^6^ cells/mL were seeded from the inlet into the microfluidic channels by gravity-driven flow. Microvessels were maintained under unidirectional gravity-driven flow for three days by replacing medium every 12 h and by keeping a difference of pressure between the inlet and outlet port of 80 Pa. After three days, the microvessels were perfused with *P. falciparum*-infected erythrocytes.

### Parasite binding assay in 3D brain microvessels

Parasite cultures were enriched for mature-stage *P. falciparum* infected erythrocytes using a MACS cell separator with LD columns (Miltenyi Biotec #130-042-901), and diluted to 10 × 10^6^ infected erythrocytes per mL in PBS. Infected erythrocytes were labeled with a membrane dye using the PKH26R Red Fluorescent Cell Linker Midi Kit (Sigma #MIDI26-1KT) according to manufacturer’s instructions. Infected erythrocytes were incubated for 30 min. at RT with either human IgG_1_ isotype control (Biolegend #403502, 0.47 mg/mL), mAb C7 (0.47 mg/mL), mAb C74 (0.40 mg/mL), or recombinant EPCR (60 μg/mL), all in PBS. Ten million infected erythrocytes per mL were perfused at 37°C for 15 min. at a flow rate of 10 μL/min using a syringe infusion pump (KD Scientific #KDS220), followed by a 10-min. wash with PBS at the same flow rate. HBMEC microvessels were fixed in 4% PFA for 20 min. followed by two 10-min. washes in PBS, and stained with DAPI (8 μg/mL) for 30 min. Each 3D microvessel device was used once for each experimental condition.

### Numerical simulation of wall shear stress rates

The flow characteristics of 3D brain microvessels during perfusion with infected erythrocytes were simulated using COMSOL Multiphysics software. Flow in the microvessel network (diameter 120 µm) was assumed to be laminar, and the stationary solver for laminar flow was used with predefined Navier-Stokes equations. Due to the low hematocrit (<0.1%) used during perfusion, flow was assumed to be Newtonian, and wall shear stress rates were calculated based on fluid viscosity of water or culture medium at 37°C (viscosity of 6.922 × 10^-4^ Pa s and density of 993.3 kg/m^3^). The inlet boundary conditions were defined for the perfusion flow rate of 10 μL/min, and the outlet boundary conditions were set at zero pressure.

### Parasite binding quantification

For each device, four edges of the 13-by-13 grid were imaged. A Zeiss LSM 980 AiryScan2 microscope with 10× NA 0.3 objective was used to image sequestered infected erythrocytes labeled with PKH26 membrane dye (laser 555 nm) and DAPI-stained parasite and HBMEC nuclei (405 nm laser). DAPI staining was used as control to confirm that the coverage of HBMECs in the microvessels was uniform (**Figure S6**). Images were acquired at a 3-μm Z-step size, and projection images of the bottom of the vessel were produced from Z-stacks using Fiji (ImageJ v1.52b) software. The percentage of parasite binding to the HBMECs was determined by thresholding the area occupied by labeled infected erythrocytes relative to a standard rectangular area of interest for 12 different of wall shear stress rates along the edges of the device. Within each edge channel, one standard region of interest of 300.64 μm × 73.84 μm was used to determine parasite binding per area. To avoid flow artifacts, parasite binding was only measured in the center of the channel where the flow is laminar and fully developed, avoiding junctions between branches. Entry and exit regions were excluded, as flow is not fully developed in these regions. Each edge was considered a technical replicate for each device, while each device was considered an independent biological replicate. To account for small differences in vessel diameter that can alter the estimated shear stress, statistical analysis of infected erythrocyte cytoadhesion under flow was performed by binning adjacent regions into shear stress increments of ∼0.5 dyn/cm^2^. To account for variability between experiments, each condition was measured in at least five different devices.

### Immunofluorescence microscopy of 3D microvessels

Immunofluorescence assays were performed using gravity flow conditions. Fixed 3D brain microvessels after perfusion were incubated in Background Buster (Innovex #NB306) for 30 min. and blocking buffer (2% bovine serum albumin, 0.1% Triton-X in PBS) for 1 hour. Blocked microvessels were stored at 4°C with mouse anti-VE-cadherin primary antibody (Abcam #ab33168), diluted 1:100 in blocking buffer. Microvessels were washed six times for 10 min. with PBS and incubated for 1 hour at RT with goat anti-mouse Alexa Fluor 647 secondary antibody, diluted 1:250 in blocking buffer (Thermo #A21235). After six 10-min. washes with PBS, vessels were imaged using a Zeiss LSM 980 confocal microscope with 20× object NA 0.8 using a 1-μm Z-step. The 3D rendering of a microvessel cross-section shown in **Figure 2** was obtained using Zeiss ZEN 3.3 and arivis Vision4D software.

### Crystallization and structure solving

Initial crystallization screening was performed by sitting-drop vapor-diffusion using the MCSG Crystallization Suite (Anatrace) using an NT8 drop setter (Formulatrix). Poorly formed crystals grew in MCSG3 well D6 (0.1 M MES, pH 6.0, 0.2 M Zn acetate, 10% PEG 8K). Crystals were optimized using the Additive Screen HT (Hampton Research). Well diffracting crystals were grown via hanging-drop vapor-diffusion using the same condition with the addition of 5% 1-propanol and were frozen with 30% glycerol as a cryoprotectant. Diffraction data were collected at Advanced Light Source beamline 5.0.2 at 12.731keV. The dataset was processed using XDS [55] and data reduction was performed using AIMLESS in CCP4 [56] to a resolution of 2.68 Å. Initial phases were solved by molecular replacement using Phaser in Phenix [57] with a search model of HB3VAR03 (PDB ID 4V3D) and mAb 258259 (PDB ID 6WTV) divided into Fv and CH1 domains. Model building was completed using Coot [58] and refinement was performed in Phenix. Additional model refinement was done using ISOLDE [59] in ChimeraX [60]. The final refinement was performed using PDB-REDO server [61]. Data collection and refinement statistics are summarized in **Table S3**.

### Cryo-EM sample preparation and data collection

C7 and C74 Fabs were complexed with the three-domain (3D) protein of IT4VAR22 at 3:1 molar ratio and incubated for 15 min. at RT. The complexes were separated by unbound Fabs and antigen by size exclusion chromatography with a Superdex 200 Increase 10/300 GL column equilibrated with Tris Buffer Saline. Fractions corresponding to the complexes were pooled and concentrated to 0.5 mg/mL. For cryo-EM, 3 µL of complexes were applied to UltrAufoil holey gold grids, and plunge frozen with a Vitrobot MarkIV (Thermo). C7 Fab was complexed with HB3VAR03 at 1.2:1 molar ratio and incubated for 20 min. at RT. Three µL of the complex sample at 0.4 mg/mL concentration was applied to UltraAufoil holey grids and plunge frozen with a Vitrobot MarkIV (Thermo).

For both C7 Fab: IT4VAR22 3D and C74 Fab:IT4VAR22 3D complexes, cryo-EM data were collected on a 200 kEV Glacios (Thermo) with Falcon IV direct electron detector. Movies were collected at a nominal magnification of 190,000×, resulting in 0.725 Å pixel size. The defocus range of -0.8 to -2.0 µm was applied for C7 Fab:IT4VAR22 3D data collection, and -0.6 to -2.2 µm was applied for C74 Fab: IT4VAR22 3D data collection. Movies with a total of 40 electron-event representation fractions were collected at a total dose of 46.77 e/Å2 and 51.33 e/Å2 for C7 Fab complex and C74 Fab complex, respectively. EPU software was used for automated data collection. C7 Fab: HB3VAR03 3D data were collected on a 200 kEV Talos Arctica (Thermo) with Gatan K2 direct electron detector. The defocus range applied was -1 to -2 µm. A total of 1850 movies were collected at a total dose of 43.5 e/Å^2^.

### Single particle Cryo-EM data processing

The C7 Fab: HB3VAR03 3D movies were aligned and dose-weighted using MotionCor2 [62]. The data processing was performed in CryoSPARC v4.1 [63]. Initial 2D templates were generated through multiple iterations of 2D classification. Using the 2D templates, particles were re-extracted using template picker and particle extraction jobs. From 145,000 particles, several iterations of 2D classification were run to screen good particles. Particles from the best 2D classes were selected to create 3D *ab-initio* maps. The best map was refined by heterogenous refinement followed by non-uniform refinement. The overall resolution of the map corresponds to ∼6 Å based on gold-standard Fourier shell correlation (GSFSC).

A total of 5187 micrograph movies were collected for C7 Fab complex data and 4628 micrograph movies were collected for C74 Fab dataset. The raw frames were dose-weighted on the fly during collection by CryoSPARC Live [63] and CTFs were estimated correspondingly. Initial particle picking was performed by Blob Picker tool in CryoSPARC v4.1 and initial 2D templates were generated through multiple iterations of 2D classification. Templates of good classes corresponding to different views of the complexes were selected as template for template picker job and a total of 1,580,114 particles from C7 Fab complex data, and 1,065,187 particles from C74 Fab complex data were re-extracted. Several iterations of 2D classifications were done to remove lower quality particles. The best 2D classes representing only the CIDRα1.7 and Fab were selected and rebalanced with 0.7 rebalance factor to distribute the particles across superclasses corresponding to different views of the complex in C7 Fab complex particle stack and a rebalance factor of 0.5 was used in the C74 Fab complex particle stack. The classes where a DBLβ3 domain was observed were discarded to aide with proper alignment in 3D classification. The best classes after rebalancing were used to generate ab-initio models with initial resolution of 8Å for the alignment of the C7 Fab complex particle, and initial and final batch size of 300 and 600 were set while 6Å initial resolution with initial and final mini-batch of 300 and 600 were used for the C74 Fab complex. The classes were then heterogeneously refined with initial resolution of 8Å. The best class with 87,881 particles of the C7 Fab complex and 207,289 particles of the C74 Fab complex was selected for non-uniform refinement. After the first round of non-uniform refinement, global CTF (beam-tilt) refinement and local CTF refinement, followed by non-uniform refinement with C1 symmetry was performed. Final map sharpening was done with B factor sharpening of -102.9 Å^2^ for the C7 Fab complex and -136.1 Å^2^ was used for the C74 Fab complex. The C7 Fab complex yielded a resolution of 3.42 Å as per GSFSC. The presence of preferred orientations reduced the quality of the data, but the overall quality of the map at the antigen-antibody binding interface had a local resolution of ∼3.2 Å. The resolution of the C74 Fab complex was estimated to be 3.35 Å as per GSFSC. Local resolution for both final maps was estimated based on a Fourier shell correlation of 0.5 and shown in **Figure S8**.

### Cryo-EM structure modeling and refinement

Initial coordinates of the CIDRα1.7 domain of IT4VAR22 were generated by AlphaFold2 [64] and docked into the cryo-EM maps of the C7 and C74 Fab complexes. Heavy and light chain variable domains of C7 and C74 Fabs were generated by ABodybuilder2 [65] and then docked into the corresponding cryo-EM maps. Once the chains were merged into a single PDB file for each of the complex structure in COOT [58], initial real space refinement followed by morphing were performed in PHENIX [66]. After manual adjustments were made in COOT, final full model refinement into the Cryo-EM maps was done with RosettaRelax [67]. General structure analysis such as Solvent Accessible Surface area measurement and RMSD calculations were performed in ChimeraX [60]. Epitope interactions were identified and visualized by Ligplot v2.6 [68]. Structural figures were generated in UCSF Chimera [69] and ChimeraX [60]. Data collection, refinement statistics, and the specific PDB and EMDB entries for cryo-EM maps and structures reported in this article are available in **Table S4**.

### Molecular dynamics simulation

To characterize the interactions and dynamics of the antibody-antigen complexes, we performed molecular dynamics simulations of C7 and the respective germline reversion. As starting structures for our simulations, we used the available cryo-EM/X-ray structures, presented in this study. We performed simulations of the free variable domains (Fvs), namely C7 and the C7 germline, to characterize the effect of the mutations on the dynamic properties of the Fvs ((227 clusters, 369 clusters) – substantial increase in variability).

The free energy conformational space molecular dynamics simulation was performed based on the method previously described in [70]. The time-lagged independent component analysis was performed to obtain kinetic discretization of the sampled conformational space [71]. The molecular dynamics analysis was performed based on the analysis described in [70]. PyMOL was used to visualize the antibody structure.

## Data availability

Maps generated from the electron microscope data are deposited in the Electron Microscopy Data Bank EMD-43148, EMD-43149, EMD-43150. Atomic models have been deposited in the RCSB Protein Data Bank with PDB IDs 8VDF, 8VDG, and 8VDL. All reagents will be made available on request after completion of a Materials Transfer Agreement. Further information and requests for resources and reagents should be directed to and will be fulfilled by the lead contact, Thomas Lavstsen.

## References

1. WHO, World malaria report 2023. World Health Organization, 2023.

2. Miller, L.H., et al., The pathogenic basis of malaria. Nature, 2002. 415(6872): p. 673-679.

3. Baruch, D.I., et al., Cloning the P. falciparum gene encoding PfEMP1, a malarial variant antigen and adherence receptor on the surface of parasitized human erythrocytes. Cell, 1995. 82(1): p. 77–87.

4. Smith, J.D., et al., Switches in expression of Plasmodium falciparum var genes correlate with changes in antigenic and cytoadherent phenotypes of infected erythrocytes. Cell, 1995. 82(1): p. 101–110.

5. Su, X.Z., et al., The large diverse gene family var encodes proteins involved in cytoadherence and antigenic variation of Plasmodium falciparum-infected erythrocytes. Cell, 1995. 82(1): p. 89–100.

6. Lau, C.K., et al., Structural Conservation Despite Huge Sequence Diversity Allows EPCR Binding by the PfEMP1 Family Implicated in Severe Childhood Malaria. Cell Host.Microbe, 2015. 17(1): p. 118–129.

7. Rask, T.S., et al., Plasmodium falciparum erythrocyte membrane protein 1 diversity in seven genomes--divide and conquer. PLoS Comput Biol, 2010. 6(9).

8. Storm, J., et al., Cerebral malaria is associated with differential cytoadherence to brain endothelial cells. EMBO Mol Med, 2019.

9. Mkumbaye, S.I., et al., The severity of Plasmodium falciparum infection is associated with transcript levels of var genes encoding EPCR-binding PfEMP1. Infect Immun, 2017.

10. Shabani, E., et al., Plasmodium falciparum EPCR-binding PfEMP1 expression increases with malaria disease severity and is elevated in retinopathy negative cerebral malaria. BMC Med, 2017. 15(1): p. 183.

11. Jespersen, J.S., et al., Plasmodium falciparum var genes expressed in children with severe malaria encode CIDRalpha1 domains. EMBO Mol Med, 2016. 8(8): p. 839–50.

12. Magallon-Tejada, A., et al., Cytoadhesion to gC1qR through Plasmodium falciparum Erythrocyte Membrane Protein 1 in Severe Malaria. PLoS Pathog, 2016. 12(11): p. e1006011.

13. Bertin, G.I., et al., Expression of the domain cassette 8 Plasmodium falciparum erythrocyte membrane protein 1 is associated with cerebral malaria in Benin. PLoS.One., 2013. 8(7): p. e68368.

14. Lavstsen, T., et al., Plasmodium falciparum erythrocyte membrane protein 1 domain cassettes 8 and 13 are associated with severe malaria in children. Proc.Natl.Acad.Sci.U.S.A, 2012. 109(26): p. E1791–E1800.

15. Rottmann, M., et al., Differential expression of var gene groups is associated with morbidity caused by Plasmodium falciparum infection in Tanzanian children. Infect.Immun., 2006. 74(7): p. 3904–3911.

16. Kessler, A., et al., Linking EPCR-Binding PfEMP1 to Brain Swelling in Pediatric Cerebral Malaria. Cell Host Microbe, 2017. 22(5): p. 601–614 e5.

17. Duffy, F., et al., Meta-analysis of Plasmodium falciparum var Signatures Contributing to Severe Malaria in African Children and Indian Adults. mBio, 2019. 10(2).

18. Tembo, D.L., et al., Differential PfEMP1 expression is associated with cerebral malaria pathology. PLoS Pathog, 2014. 10(12): p. e1004537.

19. Sahu, P.K., et al., Determinants of brain swelling in pediatric and adult cerebral malaria. JCI Insight, 2021. 6(18).

20. Bernabeu, M., et al., Severe adult malaria is associated with specific PfEMP1 adhesion types and high parasite biomass. Proc Natl Acad Sci U S A, 2016. 113(23): p. E3270–9.

21. Turner, L., et al., Severe malaria is associated with parasite binding to endothelial protein C receptor. Nature, 2013. 498(7455): p. 502-505.

22. Petersen, J.E., et al., Protein C system defects inflicted by the malaria parasite protein PfEMP1 can be overcome by a soluble EPCR variant. Thromb.Haemost., 2015. 114(5): p. 1038–1048.

23. Gillrie, M.R., et al., Diverse functional outcomes of Plasmodium falciparum ligation of EPCR: potential implications for malarial pathogenesis. Cell Microbiol., 2015. 17(12): p. 1883–1899.

24. Mosnier, L.O. and T. Lavstsen, The role of EPCR in the pathogenesis of severe malaria. Thromb Res, 2016. 141 Suppl 2: p. S46–9.

25. Obeng-Adjei, N., et al., Longitudinal analysis of naturally acquired PfEMP1 CIDR domain variant antibodies identifies associations with malaria protection. JCI Insight, 2020. 5(12).

26. Rambhatla, J.S., et al., Acquisition of Antibodies Against Endothelial Protein C Receptor-Binding Domains of Plasmodium falciparum Erythrocyte Membrane Protein 1 in Children with Severe Malaria. J Infect Dis, 2019. 219(5): p. 808–818.

27. Turner, L., et al., IgG antibodies to endothelial protein C receptor-binding cysteine-rich interdomain region domains of Plasmodium falciparum erythrocyte membrane protein 1 are acquired early in life in individuals exposed to malaria. Infect.Immun., 2015. 83(8): p. 3096–3103.

28. Tewey, M.A., et al., Natural immunity to malaria preferentially targets the endothelial protein C receptor-binding regions of PfEMP1s. mSphere, 2023. 8(5): p. e0045123.

29. Rask, T.S., et al., Plasmodium falciparum erythrocyte membrane protein 1 diversity in seven genomes--divide and conquer. PLoS.Comput.Biol., 2010. 6(9).

30. Rajan Raghavan, S.S., et al., Endothelial protein C receptor binding induces conformational changes to severe malaria-associated group A PfEMP1. Structure, 2023.

31. Lau, C.K., et al., Structural conservation despite huge sequence diversity allows EPCR binding by the PfEMP1 family implicated in severe childhood malaria. Cell Host Microbe, 2015. 17(1): p. 118–29.

32. Bernabeu, M., et al., Binding Heterogeneity of Plasmodium falciparum to Engineered 3D Brain Microvessels Is Mediated by EPCR and ICAM-1. mBio, 2019. 10(3).

33. Lipowsky, H.H., Microvascular rheology and hemodynamics. Microcirculation, 2005. 12(1): p. 5–15.

34. Leech, J.H., et al., Identification of a strain-specific malarial antigen exposed on the surface of Plasmodium falciparum-infected erythrocytes. J Exp Med, 1984. 159(6): p. 1567–75.

35. Marsh, K. and R.J. Howard, Antigens induced on erythrocytes by P. falciparum: expression of diverse and conserved determinants. Science, 1986. 231(4734): p. 150-3.

36. Udeinya, I.J., et al., Plasmodium falciparum strain-specific antibody blocks binding of infected erythrocytes to amelanotic melanoma cells. Nature, 1983. 303(5916): p. 429-31.

37. Howard, R.J., et al., Two approximately 300 kilodalton Plasmodium falciparum proteins at the surface membrane of infected erythrocytes. Mol Biochem Parasitol, 1988. 27(2-3): p. 207–23.

38. Doolan, D.L., C. Dobano, and J.K. Baird, Acquired immunity to malaria. Clin Microbiol Rev, 2009. 22(1): p. 13–36, Table of Contents.

39. Nielsen, M.A., et al., Plasmodium falciparum variant surface antigen expression varies between isolates causing severe and nonsevere malaria and is modified by acquired immunity. J Immunol, 2002. 168(7): p. 3444–50.

40. Bull, P.C., et al., Parasite antigens on the infected red cell surface are targets for naturally acquired immunity to malaria. Nat Med, 1998. 4(3): p. 358–60.

41. Marsh, K., et al., Antibodies to blood stage antigens of Plasmodium falciparum in rural Gambians and their relation to protection against infection. Trans R Soc Trop Med Hyg, 1989. 83(3): p. 293–303.

42. Bull, P.C., et al., Antibody recognition of Plasmodium falciparum erythrocyte surface antigens in Kenya: evidence for rare and prevalent variants. Infect Immun, 1999. 67(2): p. 733–9.

43. Otto, T.D., et al., Evolutionary analysis of the most polymorphic gene family in falciparum malaria. Wellcome Open Res, 2019. 4: p. 193.

44. Brazier, A.J., et al., Pathogenicity Determinants of the Human Malaria Parasite Plasmodium falciparum Have Ancient Origins. mSphere, 2017. 2(1).

45. Kilama, M., et al., Estimating the annual entomological inoculation rate for Plasmodium falciparum transmitted by Anopheles gambiae s.l. using three sampling methods in three sites in Uganda. Malar J, 2014. 13: p. 111.

46. Kamya, M.R., et al., Malaria transmission, infection, and disease at three sites with varied transmission intensity in Uganda: implications for malaria control. Am J Trop Med Hyg, 2015. 92(5): p. 903–12.

47. Azasi, Y., et al., Infected erythrocytes expressing DC13 PfEMP1 differ from recombinant proteins in EPCR-binding function. Proc Natl Acad Sci U S A, 2018. 115(5): p. 1063–1068.

48. Gonzales, S.J., et al., Naturally Acquired Humoral Immunity Against Plasmodium falciparum Malaria. Front Immunol, 2020. 11: p. 594653.

49. Kisalu, N.K., et al., A human monoclonal antibody prevents malaria infection by targeting a new site of vulnerability on the parasite. Nat Med, 2018. 24(4): p. 408–416.

50. Turner, L., et al., Severe malaria is associated with parasite binding to endothelial protein C receptor. Nature, 2013. 498(7455): p. 502-5.

51. Cham, G.K., et al., A semi-automated multiplex high-throughput assay for measuring IgG antibodies against Plasmodium falciparum erythrocyte membrane protein 1 (PfEMP1) domains in small volumes of plasma. Malar J, 2008. 7: p. 108.

52. Lennartz, F., et al., Structure-Guided Identification of a Family of Dual Receptor-Binding PfEMP1 that Is Associated with Cerebral Malaria. Cell Host Microbe, 2017. 21(3): p. 403–414.

53. Bachmann, A. and T. Lavstsen, Analysis of var Gene Transcript Patterns by Quantitative Real-Time PCR. Methods Mol Biol, 2022. 2470: p. 149–171.

54. Zheng, Y., et al., In vitro microvessels for the study of angiogenesis and thrombosis. Proc Natl Acad Sci U S A, 2012. 109(24): p. 9342–7.

55. Kabsch, W., Xds. Acta Crystallogr D Biol Crystallogr, 2010. 66(Pt 2): p. 125–32.

56. Evans, P.R. and G.N. Murshudov, How good are my data and what is the resolution? Acta Crystallogr D Biol Crystallogr, 2013. 69(Pt 7): p. 1204–14.

57. Liebschner, D., et al., Macromolecular structure determination using X-rays, neutrons and electrons: recent developments in Phenix. Acta Crystallogr D Struct Biol, 2019. 75(Pt 10): p. 861–877.

58. Emsley, P. and K. Cowtan, Coot: model-building tools for molecular graphics. Acta Crystallogr D Biol Crystallogr, 2004. 60(Pt 12 Pt 1): p. 2126-32.

59. Croll, T.I., ISOLDE: a physically realistic environment for model building into low-resolution electron-density maps. Acta Crystallogr D Struct Biol, 2018. 74(Pt 6): p. 519–530.

60. Pettersen, E.F., et al., UCSF ChimeraX: Structure visualization for researchers, educators, and developers. Protein Sci, 2021. 30(1): p. 70–82.

61. Joosten, R.P., et al., The PDB_REDO server for macromolecular structure model optimization. IUCrJ, 2014. 1(Pt 4): p. 213–20.

62. Zheng, S.Q., et al., MotionCor2: anisotropic correction of beam-induced motion for improved cryo-electron microscopy. Nat Methods, 2017. 14(4): p. 331–332.

63. Punjani, A., et al., cryoSPARC: algorithms for rapid unsupervised cryo-EM structure determination. Nat Methods, 2017. 14(3): p. 290–296.

64. Tunyasuvunakool, K., et al., Highly accurate protein structure prediction for the human proteome. Nature, 2021. 596(7873): p. 590-596.

65. Abanades, B., et al., ImmuneBuilder: Deep-Learning models for predicting the structures of immune proteins. Commun Biol, 2023. 6(1): p. 575.

66. Afonine, P.V., et al., Real-space refinement in PHENIX for cryo-EM and crystallography. Acta Crystallogr D Struct Biol, 2018. 74(Pt 6): p. 531–544.

67. Wang, R.Y., et al., Automated structure refinement of macromolecular assemblies from cryo-EM maps using Rosetta. Elife, 2016. 5.

68. Laskowski, R.A. and M.B. Swindells, LigPlot+: multiple ligand-protein interaction diagrams for drug discovery. J Chem Inf Model, 2011. 51(10): p. 2778–86.

69. Pettersen, E.F., et al., UCSF Chimera--a visualization system for exploratory research and analysis. J Comput Chem, 2004. 25(13): p. 1605–12.

70. Fernandez-Quintero, M.L., et al., Germline-Dependent Antibody Paratope States and Pairing Specific V(H)-V(L) Interface Dynamics. Front Immunol, 2021. 12: p. 675655.

71. Chodera, J.D. and F. Noe, Markov state models of biomolecular conformational dynamics. Curr Opin Struct Biol, 2014. 25: p. 135–44.

