## Supplementary Figures and Tables for "Broadly inhibitory antibodies against severe malaria virulence proteins"

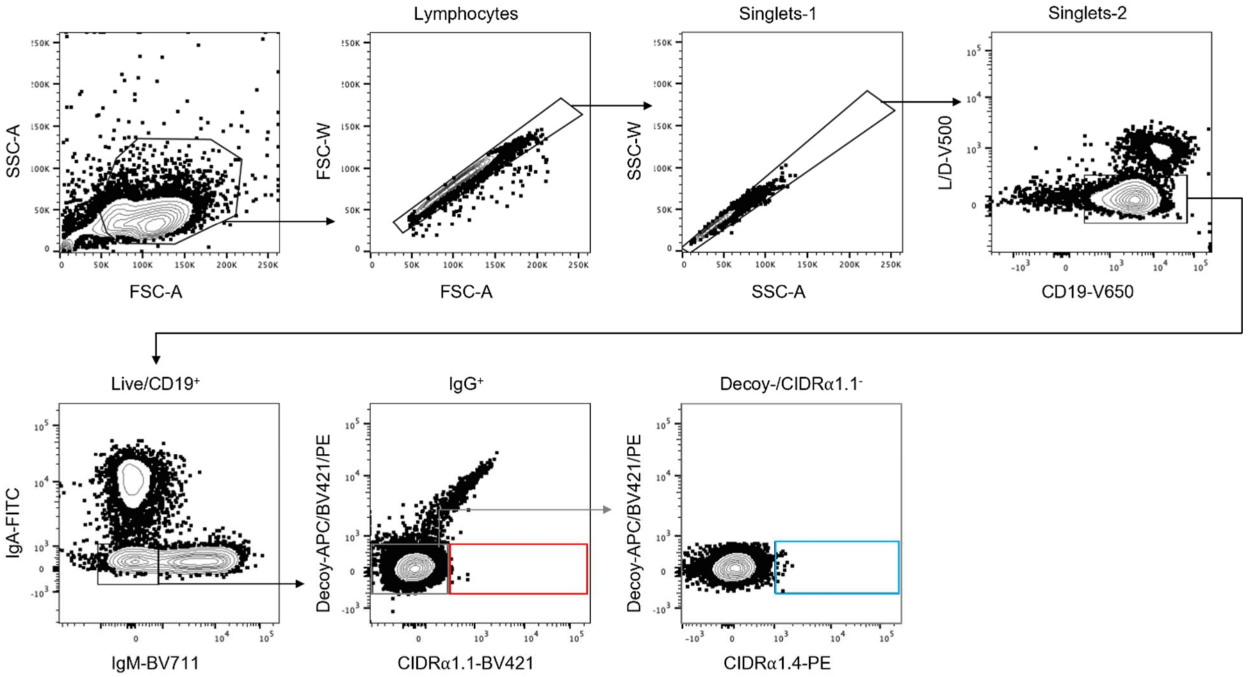


**Figure S1: Gating strategy used for sorting antigen-specific B cells.** B cells were identified by CD19 expression. Cells were negatively selected for IgG-expressing cells by excluding B cells that were IgM or IgA positive. A decoy tetramer was used to exclude non-specific cells that would bind both decoy and antigen tetramers. B cells binding to CIDRα1.1 and CIDRα1.4 tetramers were sorted separately.

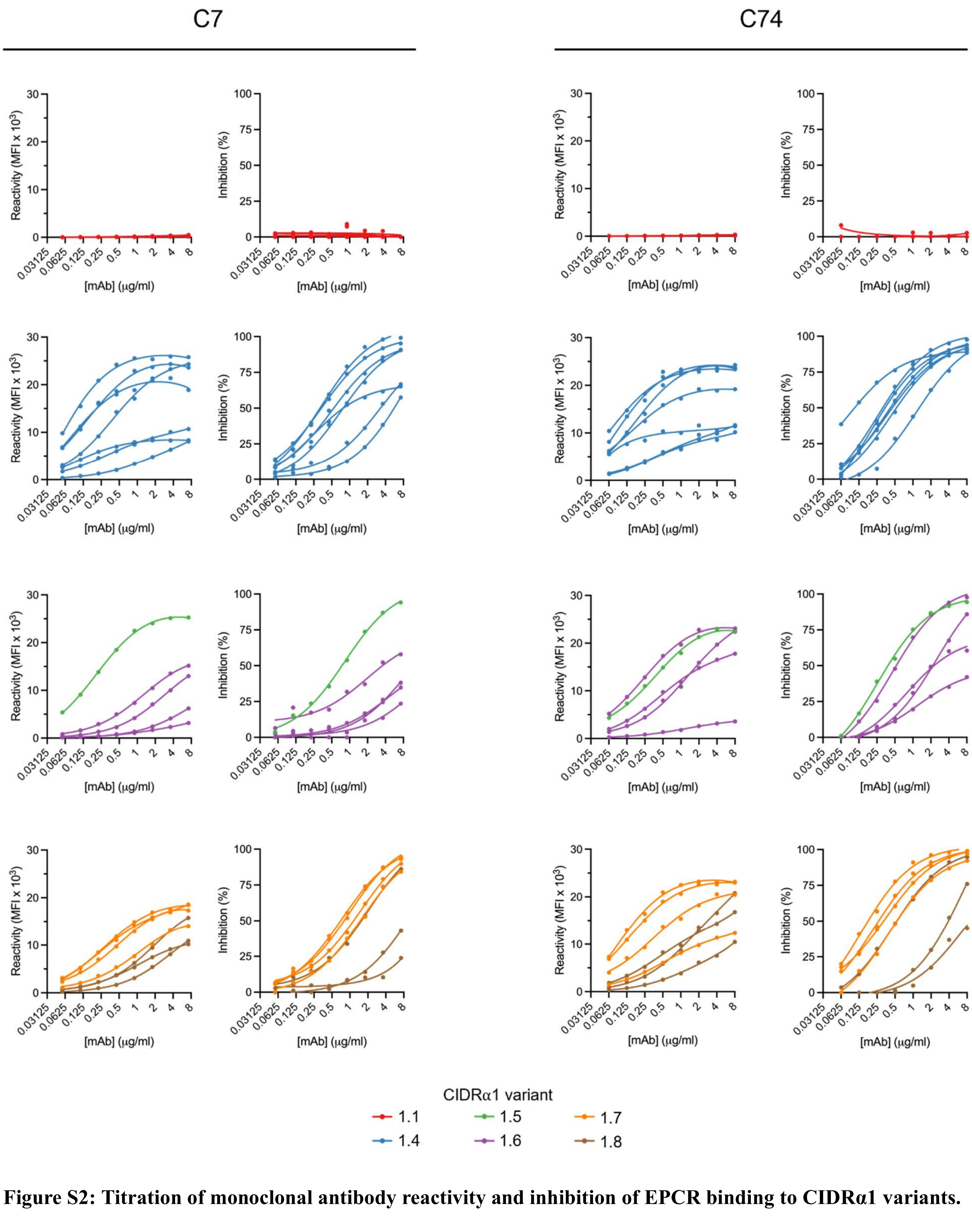


**Figure S2: Titration of monoclonal antibody reactivity and inhibition of EPCR binding to CIDRα1 variants.**

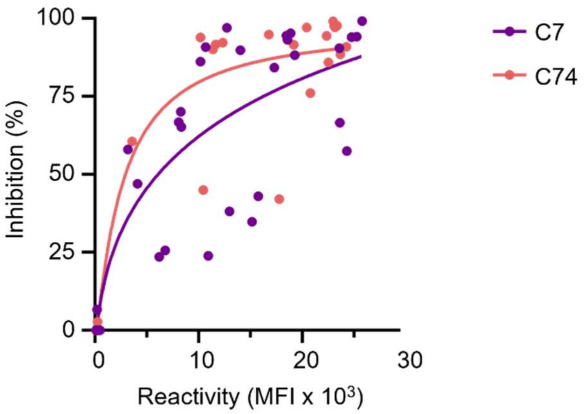


**Figure S3: Correlation of CIDRα1 binding and inhibition of EPCR binding for mAbs C7 and C74.**

**A**

| PfEMP1 variant | Protein description | Domain composition | | | | | | | | MW (kDa) | C7 R | C7 Inh | C74 R | C74 Inh |
|---|---|---|---|---|---|---|---|---|---|---|---|---|---|---|
| IT4VAR20 | full length | N | DBLα2 | CIDRα1.1 | DBLβ12 | DBLγ6 | DBLδ1 | CIDRβ1 | | 300 | neg | 0% | neg | 6% |
| IT4VAR19 | three-domain | N | DBLα2 | CIDRα1.1 | DBLβ12 | | | | | 142 | neg | 0% | neg | 4% |
| HB3VAR03 | full length | N | DBLα1.7 | CIDRα1.4 | DBLβ3 | DBLγ12 | DBLδ5 | CIDRβ3 | DBLβ7 | 330 | pos | 88% | pos | 71% |
| IT4VAR07 | N-terminal domain complex | N | DBLα1.7 | CIDRα1.4 | | | | | | 86 | pos | 91% | pos | 92% |
| IT4VAR22 | three-domain | N | DBLα1.4 | CIDRα1.7 | DBLβ3 | | | | | 138 | pos | 95% | pos | 95% |

**B**

| PfEMP1 variant | Protein description | Domain composition | | | | C7 Fab - FAB2G probe at 10 μg/mL kon [M-1s-1] | C7 Fab koff [s-1] | C7 Fab KD [nM] | C74 Fab - FAB2G probe at 10 μg/mL kon [M-1s-1] | C74 Fab koff [s-1] | C74 Fab KD [nM] |
|---|---|---|---|---|---|---|---|---|---|---|---|
| IT4VAR20 | single CIDRα1 | | | CIDRα1.1 | | No binding | | | No binding | | |
| HB3VAR03 | single CIDRα1 | | | CIDRα1.4 | | 2.24E+06 | 2.90E-04 | 0.1 | 1.56E+05 | 4.39E-04 | 2.8 |
| | three-domain | N | DBLα1.7 | CIDRα1.4 | DBLβ3 | - | - | | - | - | |
| IT4VAR07 | N-terminal domain complex | N | DBLα1.7 | CIDRα1.4 | | 3.50E+05 | 7.44E-04 | 2.1 | 3.26E+05 | 1.39E-04 | 0.4 |
| 3D7_PFD1235w | single CIDRα1 | | | CIDRα1.6 | | 8.23E+04 | 3.92E-04 | 4.8 | 1.39E+05 | 4.12E-04 | 3.0 |
| IT4VAR22 | single CIDRa1 | | | CIDRα1.7 | | 2.76E+05 | 8.91E-04 | 3.2 | 2.66E+05 | 7.91E-04 | 3.0 |
| | three-domain | N | DBLα1.4 | CIDRα1.7 | DBLβ3 | 2.84E+05 | 2.58E-04 | 0.9 | 5.26E+04 | 5.54E-04 | 10.5 |
| GA029 | single CIDRa1 | | | CIDRα1.8 | | 1.82E+04 | 3.21E-04 | 17.7 | 1.39E+04 | 3.61E-04 | 26.0 |

**C**

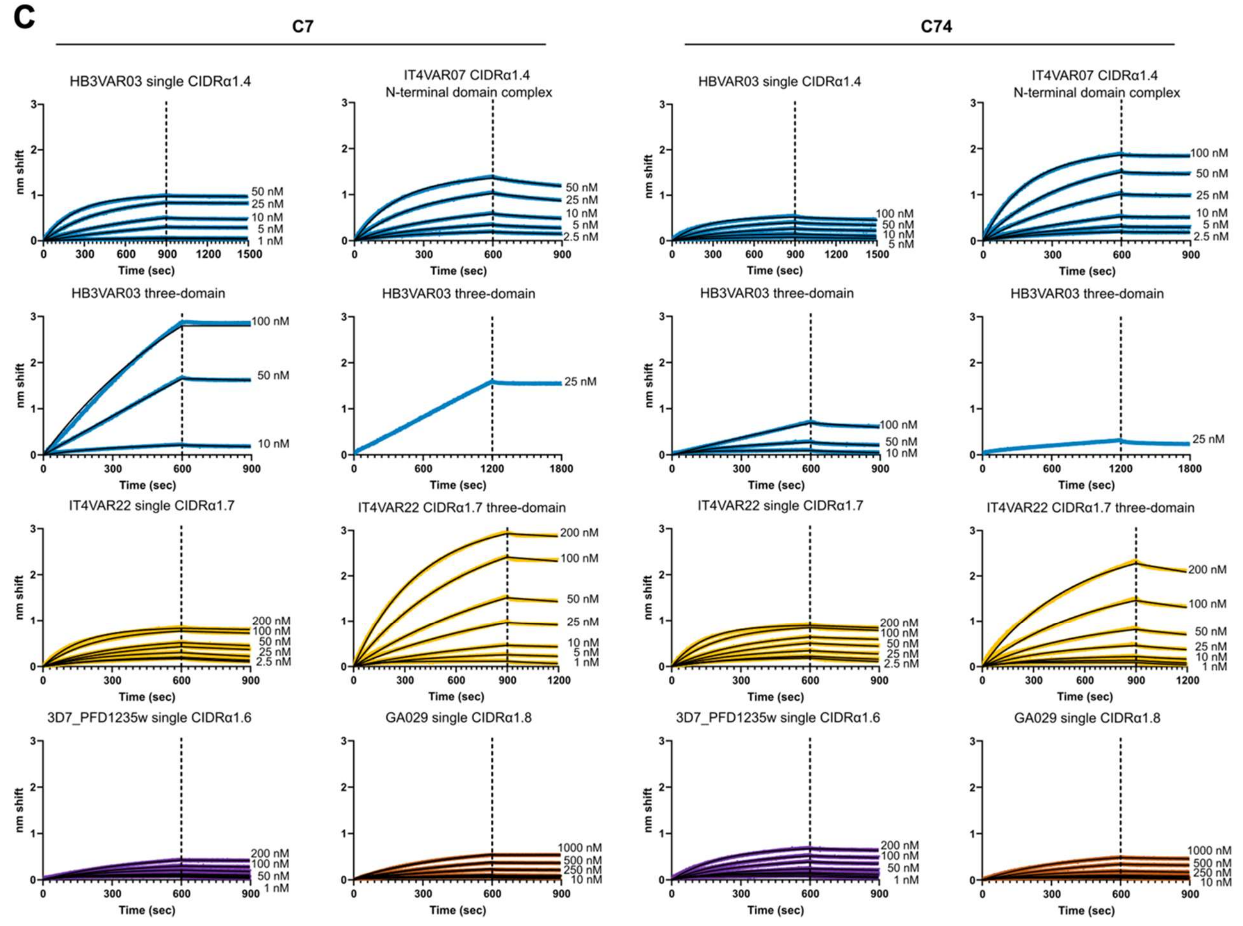


**Figure S4: Antibody binding to and inhibition of recombinant PfEMP1. A)** Binding (R) of monoclonal antibodies to recombinant PfEMP1 proteins and inhibition (Inh) of PfEMP1-EPCR binding in ELISA. Positive (pos) antibody reactivity: OD > 1.5; negative (neg): OD < 0.1. Inhibition was calculated as the percentage reduction in ELISA OD of EPCR binding to PfEMP1 protein following antibody pre-incubation as compared to a no-antibody control. N, N-terminal segment. **B)** Antibody binding kinetics to PfEMP1 proteins using biolayer interferometry (BLI). The HB3VAR03 three-domain protein showed binding to both C7 and C74 Fab but no well-fitting model was able to be applied. **C)** Representative BLI curves used to determine binding affinity to CIDRα1 domains. The raw data is colored and the 1:1 binding model fitted data is shown by the black lines. Dashed line indicates the change from association to dissociation. Concentration of the analyte is indicated next to the curves. For HB3VAR03 three-domain binding, no accurate 1:1 model could be fit as association curves remained linear even out to 1200 s.

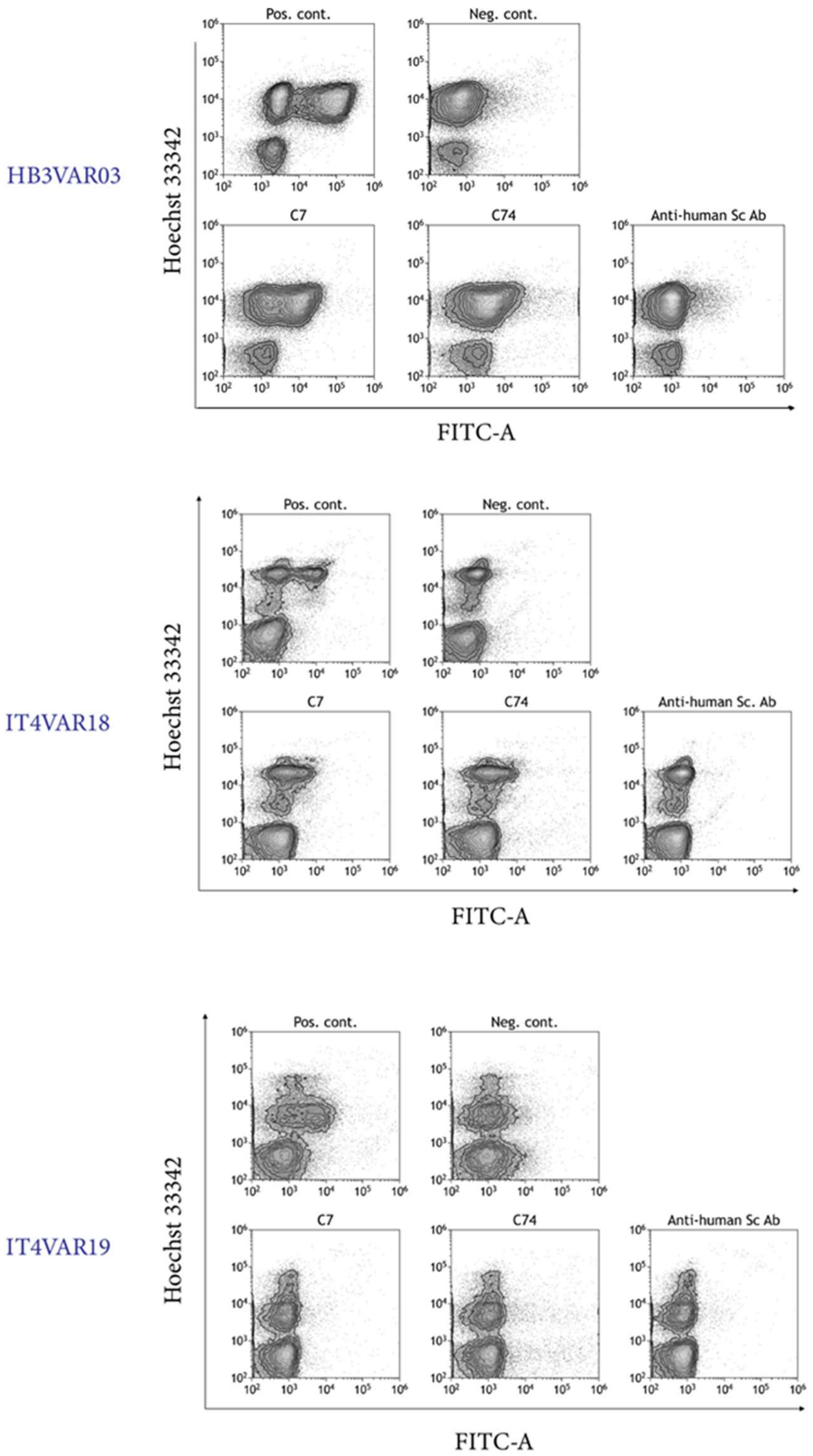


**Figure S5: C7 and C74 binding to native PfEMP1.** Flow cytometry data showing antibody (FITC-A) staining of live erythrocytes infected with HB3 and IT4 parasites expressing either CIDRα1.4 PfEMP1 (HB3VAR03), CIDRα1.6 PfEMP1 (IT4VAR18), or CIDRα1.1 PfEMP1 (IT4VAR19). Infected and uninfected erythrocytes are separated by staining of parasite DNA using Hoechst 33342. For positive controls, we used IgG from rats immunized with the full ectodomain of HB3VAR03, the CIDRα1.6 domain of IT4VAR18, or the three-domain protein of IT4VAR19. IgG from malaria-naïve individuals and FITC-conjugated goat anti-human secondary antibody were used as negative controls.

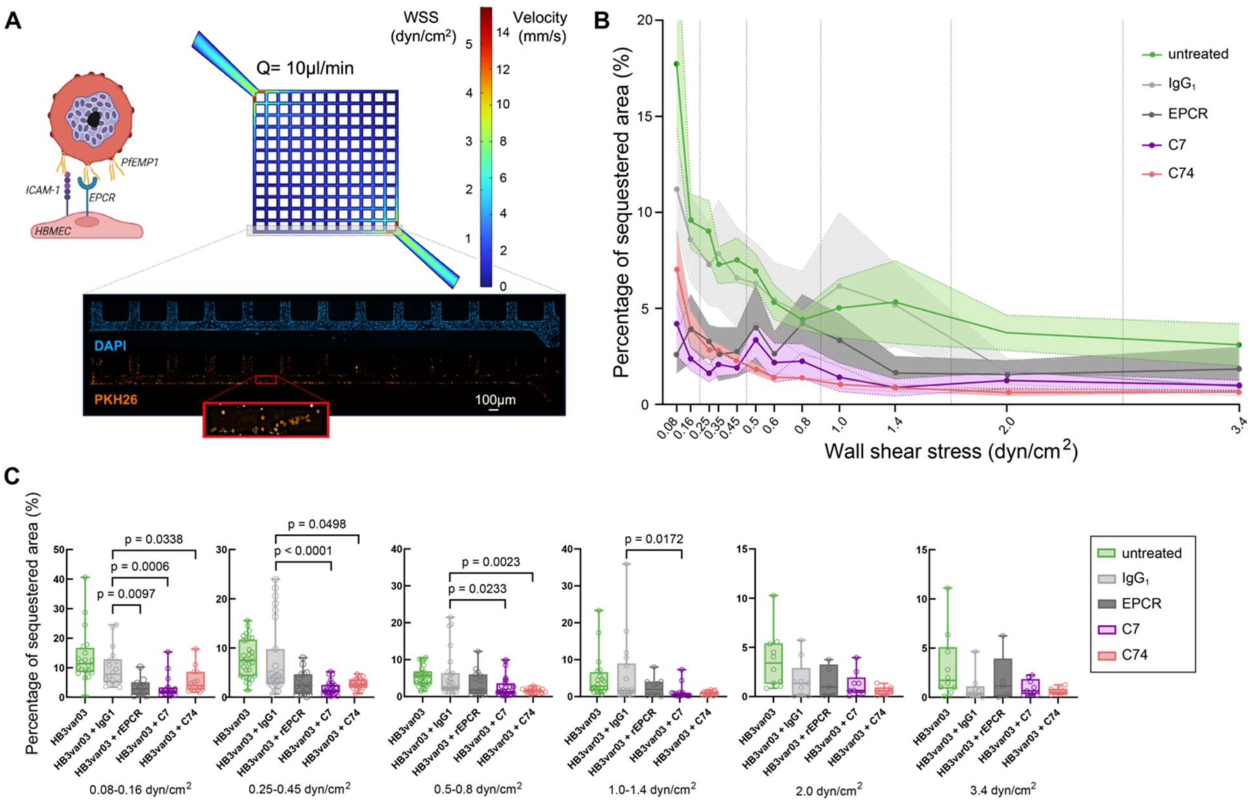


**Figure S6: Parasite binding experiments in 3D brain microvessels. A)** Top: Mid-plane flow velocity (z = 50 µm) and estimated wall shear stress (WSS; z = 0 µm) simulations in the grid geometry by COMSOL prior to collagen remodeling by human brain microvascular endothelial cells at a physiological temperature of 37°C. Bottom: Z-projection of the bottom edge of a 3D microvessel after perfusion with PKH26-labeled HB3VAR03 parasites (orange, zoom in the inset), fixation, and staining with DAPI (blue). **B)** Percentage of endothelial area occupied by sequestered *P. falciparum*-infected erythrocytes in the 3D microvessels at regions exposed to different estimated WSS rates along the edges of the device. Dots indicate the median values, and the shaded regions show the interquartile range for HB3VAR03 parasites only (green; n = 10 independent biological replicates) and after incubation with the isotype control $IgG_1$ (light gray; n = 9), recombinant EPCR (dark gray; n = 5), C7 (purple; n = 9) and C74 (red; n = 7), respectively. **C**) Percentages of sequestered area in binned regions (denoted by dotted lines in panel B) are shown in boxplots. Statistical analyses were performed for binned regions (dotted vertical lines) using a Kruskal-Wallis test, followed by comparisons between $IgG_1$ and C7, C74, or EPCR using Dunn's post-hoc test, corrected for multiple comparisons. Horizontal lines indicate medians, whiskers represent the range from the minimum to the maximum value.

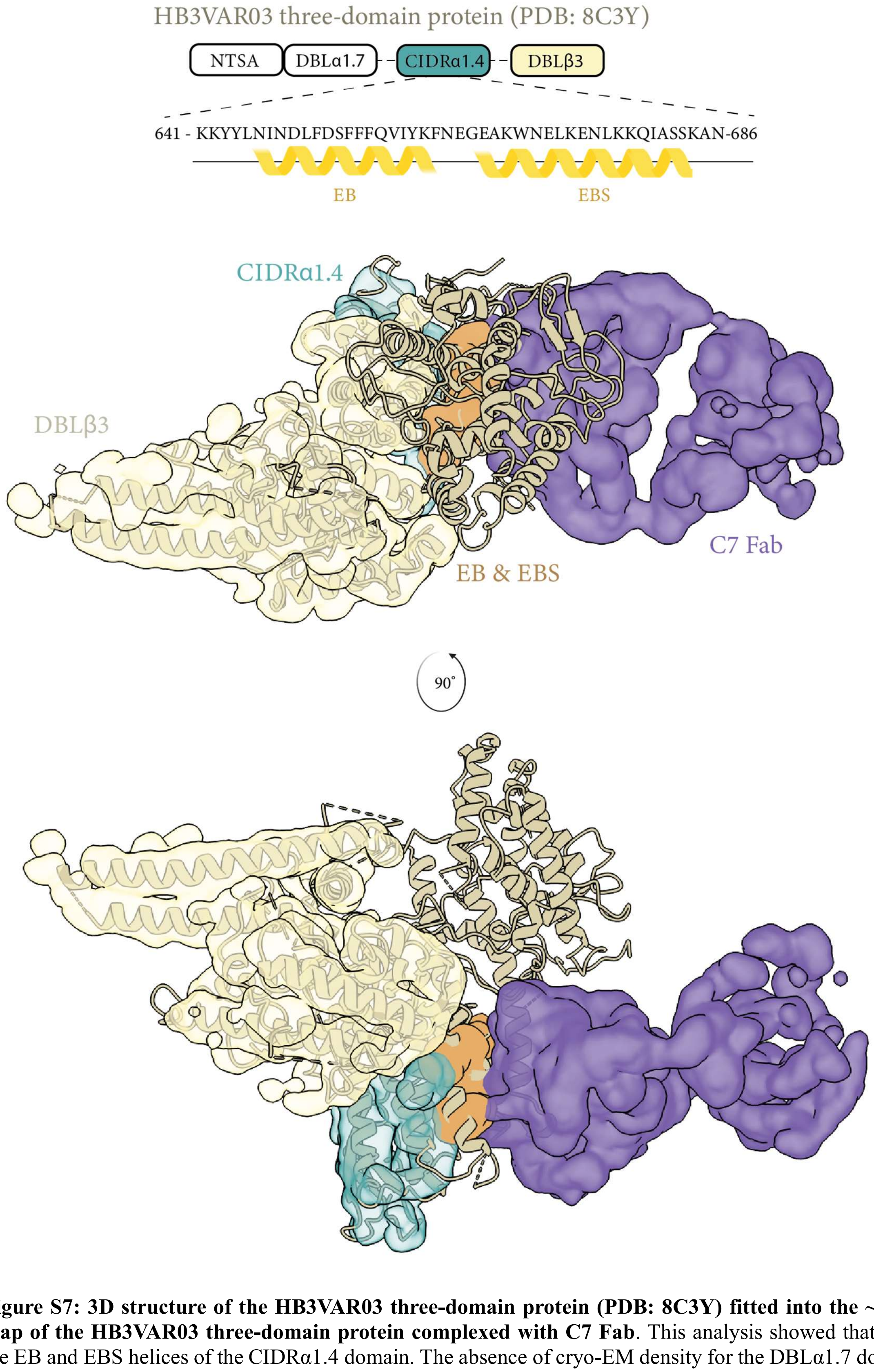


**Figure S7: 3D structure of the HB3VAR03 three-domain protein (PDB: 8C3Y) fitted into the ~6 Å cryo-EM map of the HB3VAR03 three-domain protein complexed with C7 Fab**. This analysis showed that C7 Fab binds the EB and EBS helices of the CIDRα1.4 domain. The absence of cryo-EM density for the DBLα1.7 domain suggests an induced flexibility in the domain's position following C7 binding.

IT4VAR22 three-domain protein complexed to C7 Fab

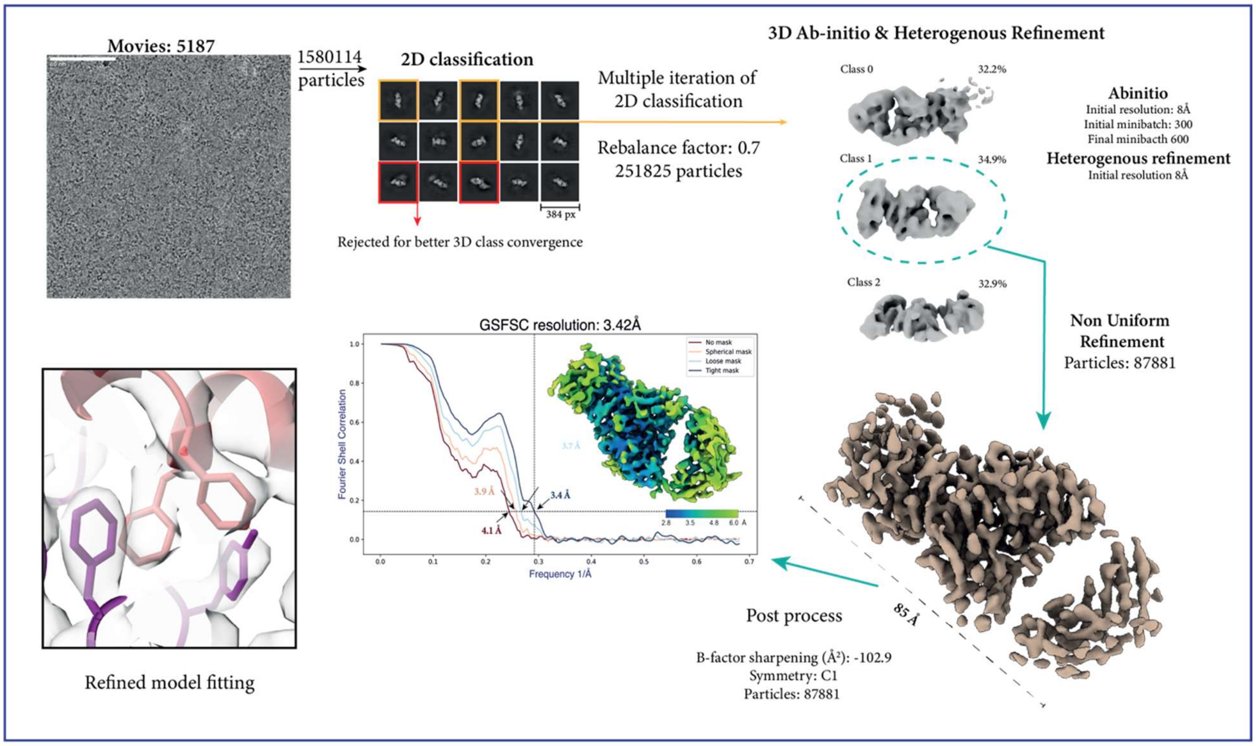


IT4VAR22 three-domain protein complexed to C74 Fab

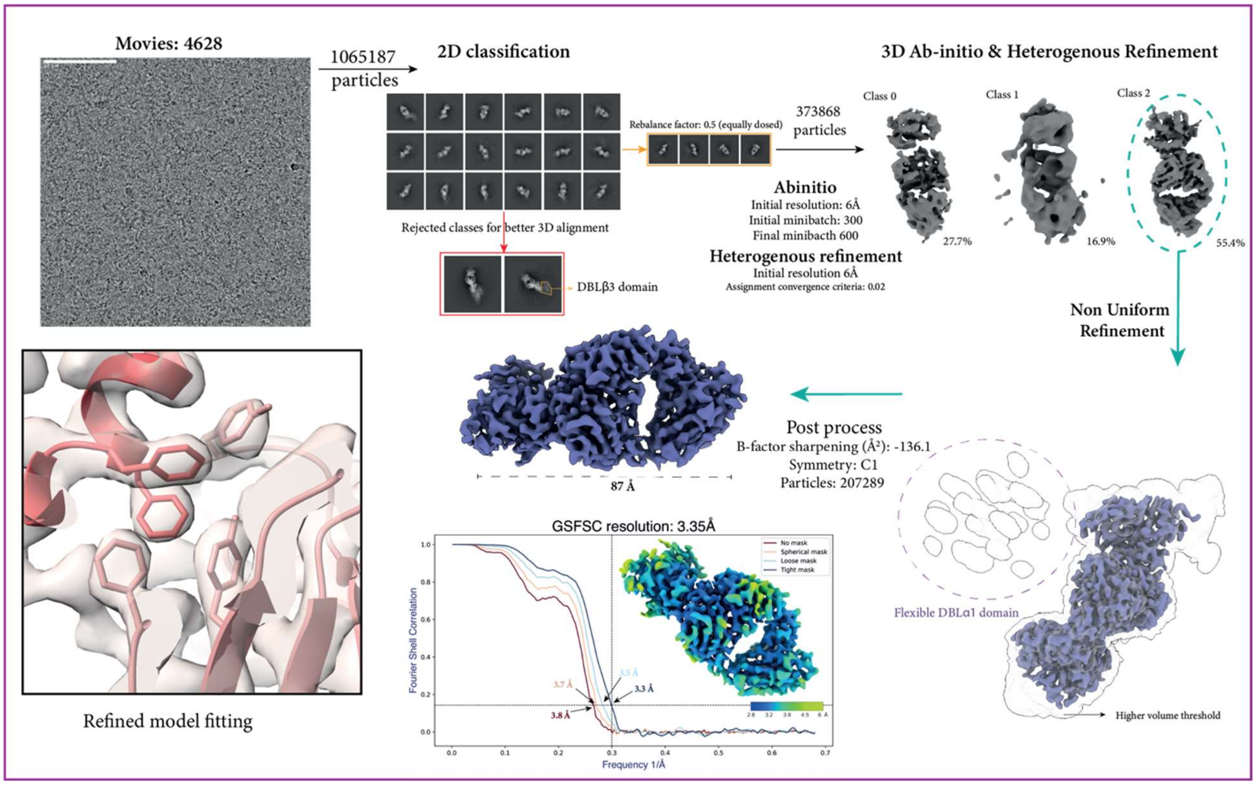

**Figure S8: Single particle cryo-EM workflow in solving IT4VAR22 three-domain protein in complex with C7 and C74 Fabs.** Cryo-EM workflow is shown, including representative micrograph of the cryo-EM data and representative 2D class averages. 2D classes in yellow boxes were selected, rebalanced to perform 3D *ab-initio* and heterogenous refinement. Red boxes contain particle stacks with the DBLα1 and DBLβ3 domains aligned, which were discarded to reduce noise in 3D alignments in both C7 and C74 data sets. Final refinement steps include non-uniform refinement followed by B-factor sharpening. The overall gold-standard Fourier shell correlation (GSFSC) represents the resolution calculated at FSC 0.143. The local resolution of the final maps was colored based on resolution calculated at FSC 0.5. The refined model was fitted into the cryo-EM map to show the quality of side chain fitting at the antigen-antibody interface. The workflow is described in more detail in the Methods section.

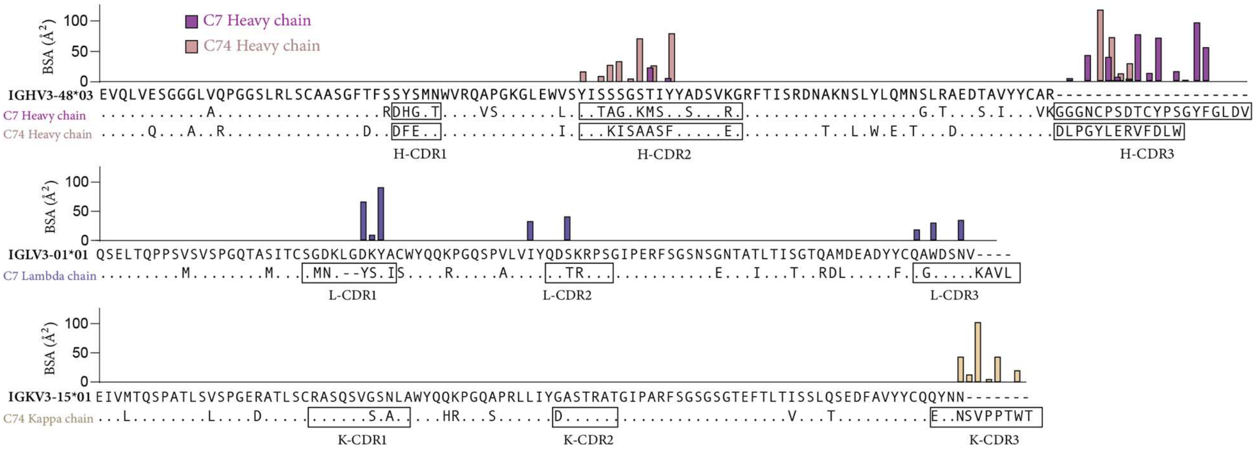


**Figure S9: Comparison of C7 and C74 mAb somatic hypermutations and buried surface area.** The buried surface area (BSA) (PDBePISA) is shown against the C7 and C74 heavy and light variable chain sequences (up to the end of the CDR3) aligned with their respective germline sequences. CDRs were defined using the Kabat numbering system.

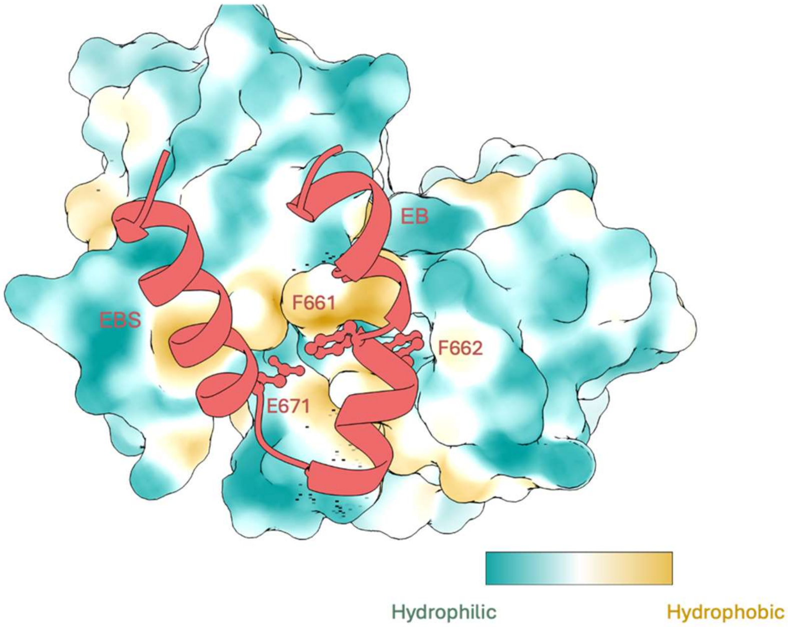


**Figure S10: Hydrophobicity of the C7 variable domain surface.** The FF motif and the glutamic acid residue of the EB and EBS helices of CIDRα1.7 (IT4VAR22; shown in red) protrude into the hydrophobic surface of the variable domain of C7 Fab, (shown in green/white/yellow).

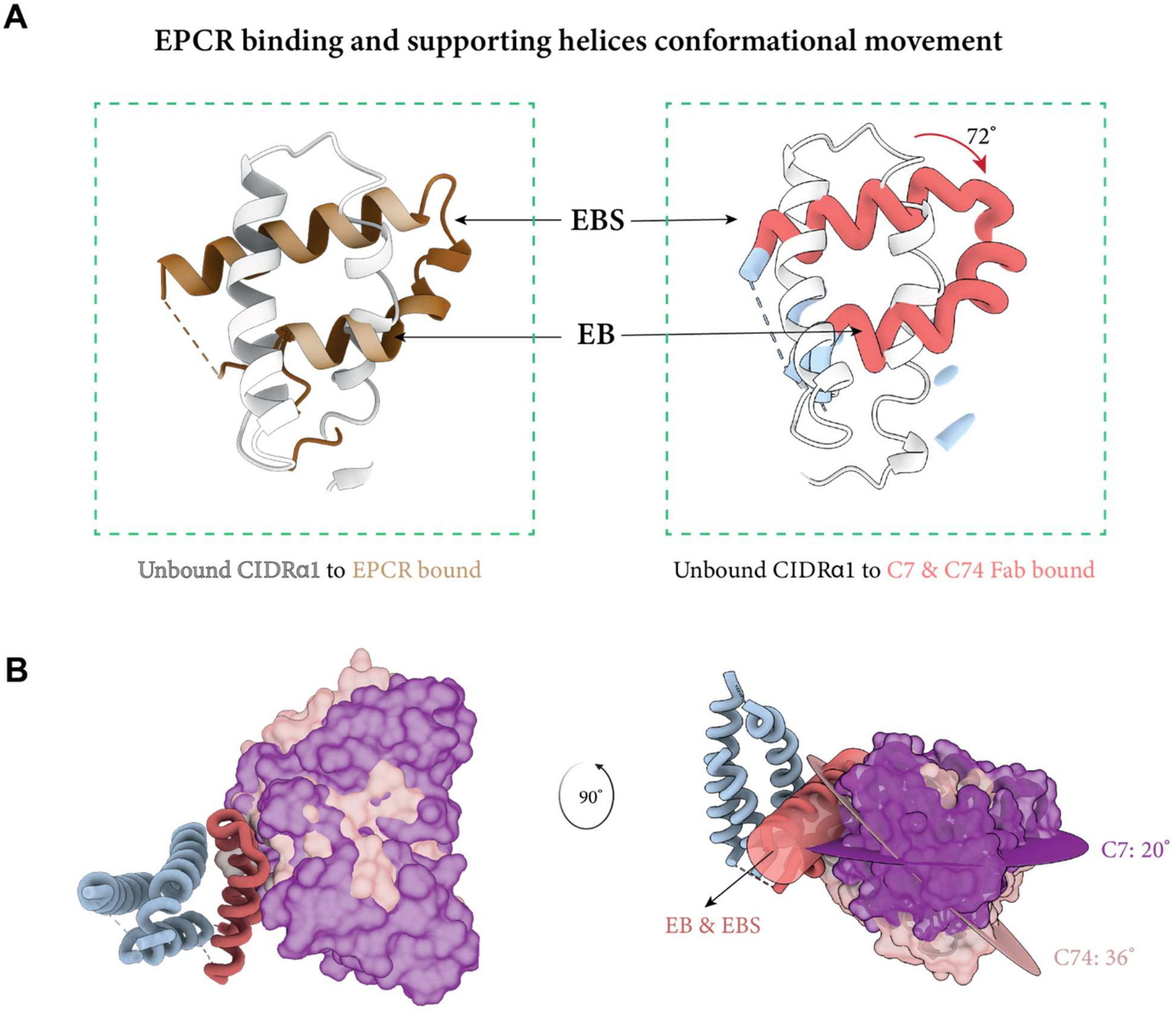


**Figure S11: Comparison of the EPCR binding site when bound and unbound by EPCR, C7, and C74. A)** Conformation of the EPCR binding (EB) and EPCR binding supporting (EBS) helices in the unbound state (white) and upon EPCR and C7 Fab binding, which induces a twist-turn conformational change (in brown and red, respectively). **B)** Depiction of the angle of approach by C7 and C74 to the axis of the EB and EBS helices of IT4VAR22 CIDRα1.7.

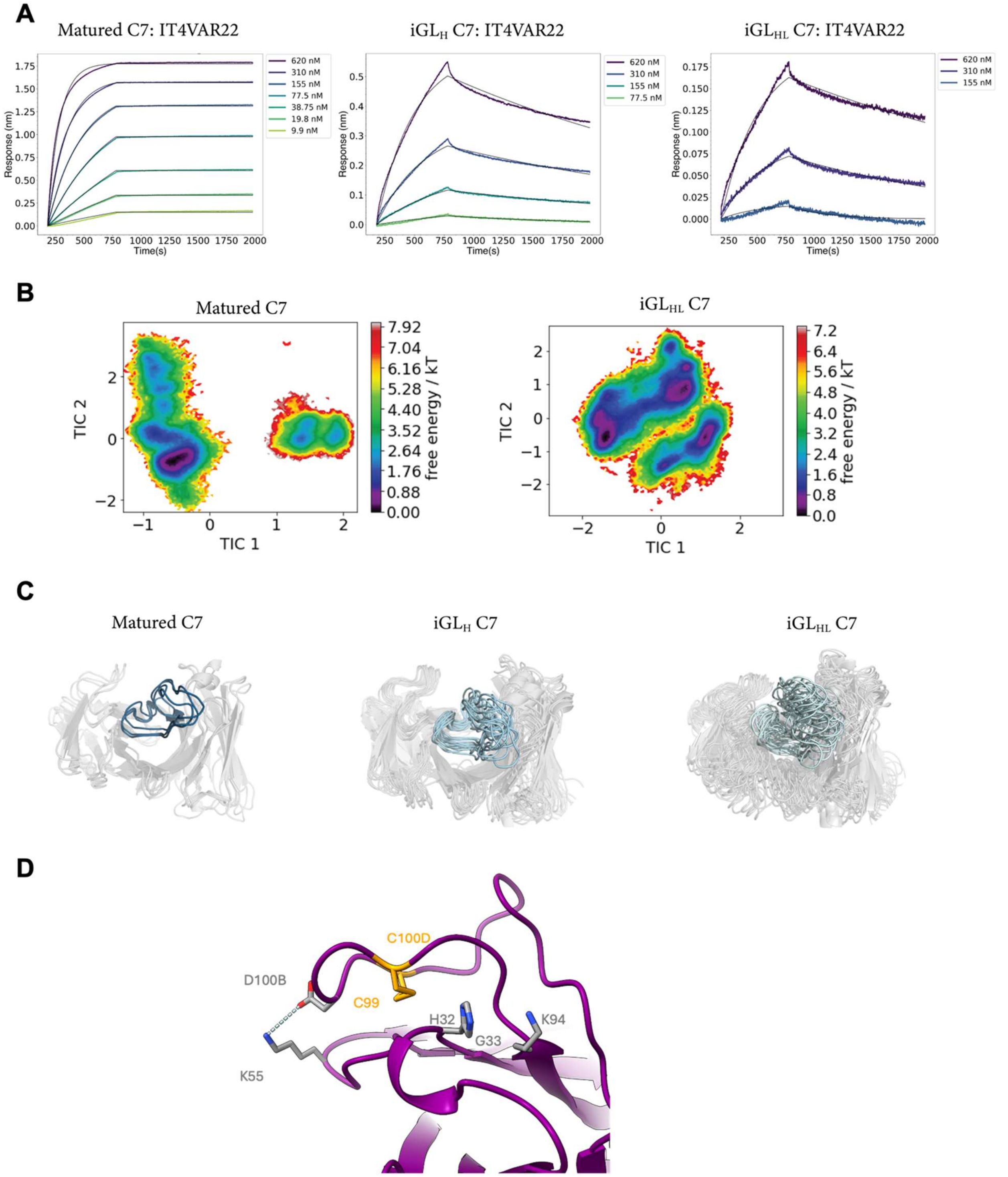


**Figure S12: Analysis of inferred germline versions of C7. A)** Biolayer interferometry analysis and comparison of matured C7, C7 with inferred germline heavy chain (iGL$_H$ C7) and C7 with inferred germline heavy and light chain (iGL$_{HL}$) binding to the IT4VAR22 three-domain protein. **B)** Free energy landscape of the H-CDR3 loop based on molecular dynamics simulations on the antigen-free matured C7 and iGL$_{HL}$ C7 in solution **C)** Conformational diversity of the H-CDR3 loop in matured C7, iGL$_H$ C7, and iGL$_{HL}$ C7. **D)** The conformational stability of the C7 H-CDR3 loop is maintained through both H-CDR1 and H-CDR2 contacts.

### Table S1: Sequence characteristics of anti-CIDRα1 monoclonal antibodies

| Donor | mAb | VH | DH | JH | HCDR3 | VH mutations (%) | VL | JL | LCDR3 | VL mutations (%) |
|---|---|---|---|---|---|---|---|---|---|---|
| 2 | C7 | V3-48 | D2-15 | J6 | CVKGGGNCPSDTCYPSGYFGLDVW | 16.0 | V3-1 | J2 | CQGWDSNVKAVLF | 13.6 |
| 2 | C62 | V3-48 | D2-2 | J6 | CVKGGGNCPSDTCYPSGYFGLDVW | 16.0 | V3-1 | J2 | CQGWDRHAKAAVF | 14.7 |
| 2 | C50 | V3-30 | D5-18 | J6 | CVRSRHGYSFTNMDVW | 12.2 | V1-51 | J1 | CGAWDDRLKIFVF | 10.5 |
| 2 | A50 | V1-69 | D2-8 | J3 | CARGAGSAGLDVW | 13.9 | V3-21 | J2 | CQVWDSGSVQVVF | 6.8 |
| 3244 | C74 | V3-48 | D3-3 | J4 | CARDLPGYLERVFDLW | 14.9 | V3-15 | J1 | CQEYKNSVPPTWTF | 9.3 |
| 3244 | A65 | V1-2 | D2-21 | J6 | CARDVVSDYYYYYMDVW | 11.5 | V4-1 | J1 | CQQYYNTPPTF | 6.7 |
| 3341 | B57 | V1-69 | D2-2 | J4 | CAKGAQTYALLYYFDSW | 15.6 | V2-8 | J2 | CSSYAYNNNLVF | 6.3 |

**Table S2: Aromatic contacts between C7, iGL$_H$ C7, iGL$_{HL}$ C7 and IT4VAR22 CIDRα1.7 predicted by molecular dynamics simulations**

| C7: CIDRα1.7 | | |
|---|---|---|
| Antigen | Antibody | Frequency |
| **E671** | Y100E | 1.0 |
| N675 | Y29 (L-CDR1) | 0.9 |
| **E671** | Y29 (L-CDR1) | 0.9 |
| **F661** | Y100I | 0.8 |
| **F662** | F100J | 0.8 |
| **F662** | Y100E | 0.8 |
| **F662** | T100C | 0.8 |
| M665 | Y100E | 0.8 |
| N654 | Y100I | 0.7 |
| E658 | Y100I | 0.7 |
| **F661** | Y100E | 0.7 |
| W674 | Y29 (L-CDR1) | 0.6 |
| **F662** | C100D | 0.5 |
| **F661** | Y29 (L-CDR1) | 0.5 |

| iGL$_H$ C7: CIDRα1.7 | | |
|---|---|---|
| Antigen | Antibody | Frequency |
| **F661** | Y100I | 1.0 |
| **E671** | Y100I | 0.9 |
| **F662** | Y100E | 0.8 |
| W674 | Y100E | 0.7 |
| **F662** | C100D | 0.6 |
| **F662** | F100J | 0.6 |
| **F662** | T100C | 0.5 |
| **F661** | Y29 (L-CDR1) | 0.5 |

| iGL$_{HL}$ C7: CIDRα1.7 | | |
|---|---|---|
| Antigen | Antibody | Frequency |
| F661 | Y100I | 0.7 |
| F662 | T100C | 0.6 |
| E671 | Y100J | 0.6 |

**Table S3: Crystallization data collection and refinement statistics**

| | HB3VAR03 CIDRα1.4 domain with C7 Fab |
|---|---|
| **Data collection** | |
| **Space group** | $P22_12_1$ |
| **Cell dimensions** | |
| ***a*, *b*, *c* (Å)** | 55.65. 55.65, 250.34 |
| ***α*, *β*, *γ* (°)** | 90, 90, 90 |
| **Resolution (Å)** | 46.34-2.68 (2.78-2.68) |
| $\boldsymbol{R}_{\mathbf{merge}}$[a] | 0.034 (0.496) |
| **<I/σ(I)>** | 11.74 (1.57) |
| $\mathbf{CC_{1/2}}$ | 0.999 (0.669) |
| **Completeness** | 99.6 (97.1) |
| **Redundancy** | 2.0 (1.99) |
| **Refinement** | |
| **Resolution (Å)** | 46.34-2.68 (2.75-2.68) |
| **No. unique reflections** | 26379 (2506) |
| $\boldsymbol{R}_{\mathbf{work}}$[b]/$\boldsymbol{R}_{\mathbf{free}}$[c] | 20.47/24.71 (31.00/32.00) |
| **No. atoms** | 4072 |
| **Protein** | 3929 |
| **Water** | 138 |
| **Ligand** | 5 |
| **B-factors (Å²)** | 9.35 |
| **Protein** | 8.73 |
| **Water** | 25.96 |
| **Ligand** | 37.80 |
| **RMS bond length (Å)** | 0.617 |
| **RMS bond angle (°)** | 0.760 |
| **Ramachadran plot statistics**[d] | |
| **Residues** | 510 |
| **Most favored region** | 95.35 |
| **Allowed region** | 4.44 |
| **Disallowed region** | 0.40 |
| **Clash score** | 8.13 |
| **PDB ID** | 8VDL |

$R_{merge} = [\sum_h\sum_i|I_h - I_{hi}|/\sum_h\sum_i I_{hi}]$ where $I_h$ is the mean of $I_{hi}$ observations of reflection *h*. Numbers in parenthesis represent highest resolution shell. [b] $R_{factor}$ and [c] $R_{free} = \sum||F_{obs}| - |F_{calc}|| / \sum|F_{obs}| \times 100$ for 95% of recorded data ($R_{factor}$) or 5% data ($R_{free}$). [d] Determined using MolProbity (10.1002/pro.3330)

**Table S4: Cryo-EM data collection, processing, model building and refinement parameters**

| | IT4VAR22 three-domain protein complexed with C7 Fab | IT4VAR22 three-domain protein complexed with C74 Fab | HB3VAR03 three-domain protein complexed with C7 Fab |
|---|---|---|---|
| **PDB** | 8VDF | 8VDG | - |
| **EMDB** | 43148 | 43149 | 43150 |
| **Microscope** | TFS Glacios | TFS Glacios | Talos Arctica |
| **Detector** | Falcon 4 | Falcon 4 | Gatan K2 |
| **Voltage** | 200kV | 200kV | 200kV |
| **Recording mode** | counting | counting | counting |
| **Total dosage (e/Å$^2$)** | 46.77 | 51.33 | 43.5 |
| **Nominal magnification** | 190000x | 190000x | 36000x |
| **Defocus range (μm)** | -0.8 to -2.0 | -0.6 to -2.2 | -1 to -2 |
| **Pixel size (Å)** | 0.725 | 0.725 | 1.15 |
| **EER number of fractions** | 40 | 40 | - |
| | **EM data processing** | | |
| **Micrograph movies collected** | 5187 | 4628 | 1850 |
| **Total particles used** | 1580114 | 1065187 | 145000 |
| **Refined particles** | 80835 | 207289 | 55000 |
| **Symmetry imposed** | C1 | C1 | C1 |
| **Map resolution (FSC 0.143) Å** | 3.42 | 3.36 | 5.2 |
| **Map sharpening B-factor (Å$^2$)** | -102.9 | -136.1 | -65.4 |
| | **Refinement and validation** | | |
| **Composition** | | | |
| Chains | 3 | 3 | |
| Atoms | 2584 | 2437 | |
| Ligands | 0 | 0 | |
| **RMSD from ideal** | | | |
| Bond length(Å) | 0.022 | 0.024 | |
| Bond angles (˚) | 1.7 | 1.71 | |
| **Clash score (all-atom)** | 0.39 | 4.9 | |

| | | |
|---|---|---|
| **Molprobity score** | 0.64 | 1.46 |
| **EMRinger score** | 2.625 | 4.25 |
| **Rotamer outlier (%)** | 0 | 0 |
| **Ramachandran plot** | | |
| Outlier (%) | 0 | 0 |
| Allowed (%) | 1.91 | 3.17 |
| Favored (%) | 98.09 | 96.83 |
| **Cβ outliers (%)** | 0 | 0 |
| **Cablam outlier (%)** | 1.99 | 2.26 |
| **Resolution estimates (Å)** | | |
| d FSC model (0/0.143/0.5) Å | 3.2/3.4/3.9 (Masked) | 3.2/3.3/3.5 (Masked) |
